# Sucrose metabolism and candidate genes during catkin fibers development in poplar

**DOI:** 10.1101/2022.05.27.493775

**Authors:** Xiong Yang, Tianyun Zhao, Pian Rao, Ning Yang, Guolei Li, Liming Jia, Xinmin An, Zhong Chen

**Affiliations:** State Key Laboratory for Efficient Production of Forest Resources, Key Laboratory of Silviculture and Conservation of the Ministry of Education, National Energy R&D Center for Non-food Biomass, Engineering Research Center for Carbon Sequestration and Sink Enhancement by Forestry and Grass of the Ministry of Education, College of Forestry, Beijing Forestry University, Beijing 100083, China; National Engineering Research Center of Tree Breeding and Ecological Restoration, College of Biological Sciences and Biotechnology, Beijing Forestry University, Beijing 100083, China

**Keywords:** Catkin fibers development, Gene expression, *Populus tomentosa*, Regulatory network, Sucrose metabolism, Transgenic lines.

## Abstract

Poplar is an important tree species for ecological protection, wood production, bioenergy and urban greening; it has been widely planted worldwide. However, the catkin fibers produced by female poplars can cause environmental pollution and safety hazards during spring. This study focused on *Populus tomentosa*, and revealed the sucrose metabolism regulatory mechanism of catkin fibers development from morphological, physiological and molecular aspects. Paraffin section suggested that poplar catkin fibers were not seed hairs and produced from the epidermal cells of funicle and placenta. Sucrose degradation via invertase and sucrose synthase played the dominant role during poplar catkin fibers development. The expression patterns revealed that sucrose metabolism-related genes played important roles during catkin fibers development. Y1H analysis indicated that there was a potential interaction between *sucrose synthase 2 (PtoSUS2)/vacuolar invertase 3 (PtoVIN3)* and MYB/ bHLH transcription factors in poplar. Finally, the two key genes, *PtoSUS2* and *PtoVIN3*, had roles in *Arabidopsis* trichome density, indicating that sucrose metabolism is important in poplar catkin fibers development. This study is not only helpful for clarifying the mechanism of sucrose regulation during trichome development in perennial woody plants, but also establishes a foundation to solve poplar catkin fibers pollution through genetic engineering methods.

**Highlight:** Sucrose degradation via invertase and sucrose synthase plays an important role in poplar catkin fibers development, and *PtoSUS2* and *PtoVIN3* are potential promising targets to solve poplar catkin fibers pollution.

## Introduction

Poplar, widely distributed and cultivated in the world, has significant economic and ecological values. Poplar is not only major contributor to wood production, but also provides resource for eco-environment construction, biomass energy, carbon sequestration and urban greening (Sannigrahi *et al*., 2010). Furthermore, poplar is a crucial model system for research on perennial woody plants due to its small genome, easy propagation and rapid growth (An *et al*., 2022). The growth rate of female poplars in the early stage is significantly faster than that of male trees, so they are more selected for production. But, in female poplars, the capsules contain seeds and catkin fibers (Fig. 1a), and begin to open as the seeds become mature (Fig. 1b). Catkin fibers cover the whole catkins (Fig. 1c, d), which is a common phenomenon in poplar (Fig. 1e, f), and the fibers can be carried over long distances with seeds on the wind (Fig. 1g, h). The long period (>40 days) during which poplar fibers are suspended in the air can cause serious environmental pollution and may pose a safety hazard due to their inflammability (Ye *et al*., 2014), although it can be used as raw materials for bioethanol (Zhang *et al*., 2018), oil super absorbents (Likon *et al*., 2013) and textile bulk insulators (Chen and Cluver, 2010). However, as its single-cell origin, the poplar catkin fiber is also specific material for studying plant cell elongation and development process, especially for tree-specific traits and perennial angiosperm development. In recent years, studies of sex differentiation and female flower development in poplar have revealed some key genes such as *type-A cytokinin response regulator* (Yang *et al*., 2021), *FERR-R* and *MSL* (Xue *et al*., 2020), *ARABIDOPSIS RESPONSE REGULATOR 17* (Müller *et al*., 2020), *AGAMOUS* and *SEEDSTICK* (Lu *et al*., 2019), and *LEAFY* (Klocko *et al*., 2016). However, the molecular basis of catkin fibers formation and development in poplar is still unclear and key genes involved in this process are not found yet. Therefore, analyzing the development process of catkin fibers is helpful not only for understanding this progress in Salicaceae plants, but also for improving the urban environment.

**Figure 1.**
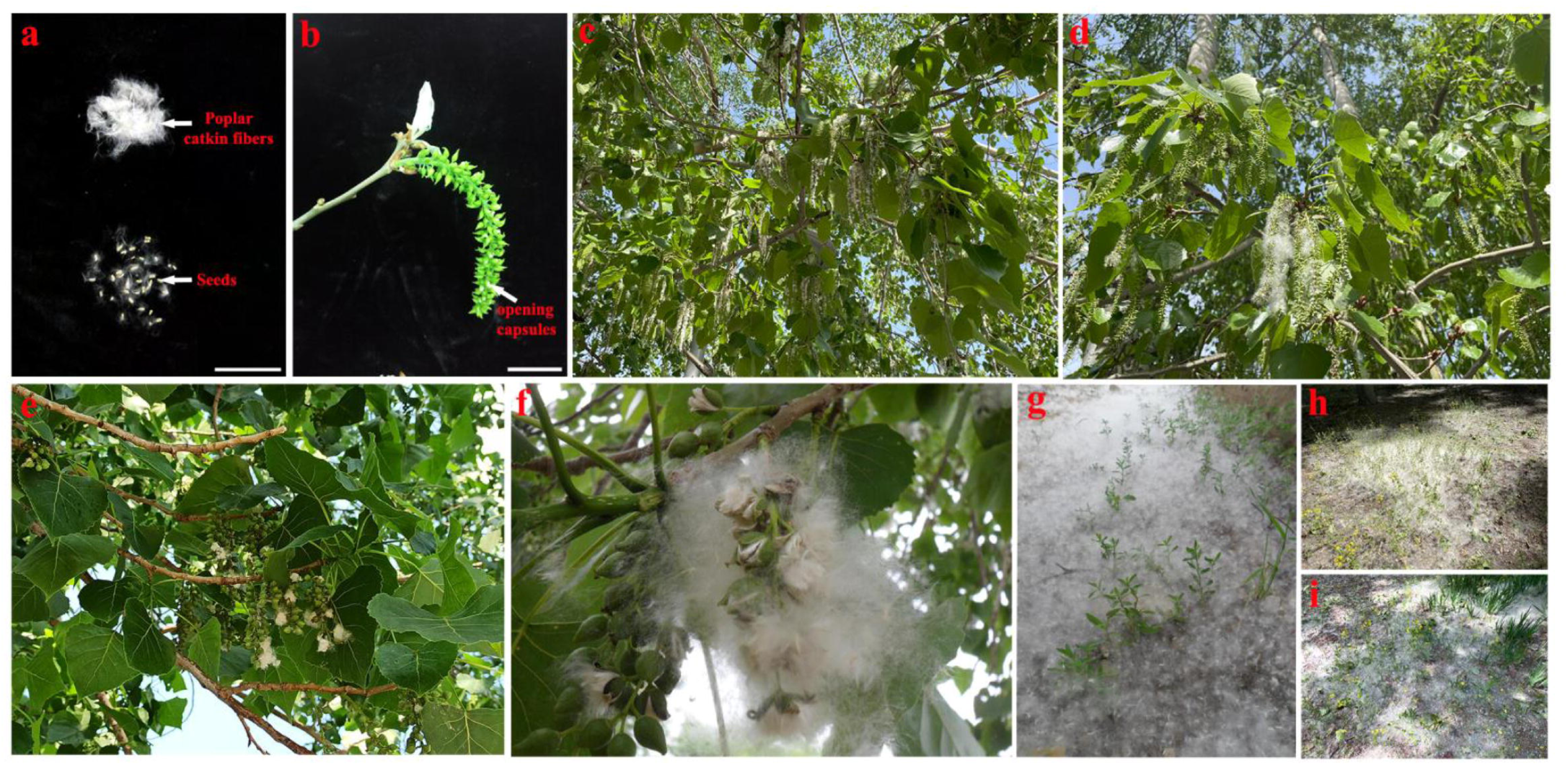
Morphological characteristics of capsules and catkin fibers in poplar. a-d. Characteristics of *Populus tomentosa.*. **e, f** Other poplars. **g-h** Environmental pollution caused by poplar catkin fibers. Scale bar: **a** = 1 cm, **b** = 2 cm.

Trichomes, which originate from epidermal cells, are distributed on the surfaces of plants and protect them from biotic and abiotic stresses (Pattanaik *et al*., 2014). Many previous studies of trichomes have been conducted in *Arabidopsis thaliana* and cotton, focusing on epidermal hairs and cotton fibers, respectively (Naoumkina, 2018, Grebe, 2012). According to morphological characters, trichome development can be divided into three stages: initiation, extension and maturation. An additional stage, secondary cell wall synthesis, is presumed to occur between extension and maturation in cotton fibers (Naoumkina, 2018, Grebe, 2012). Similar regulatory mechanisms were observed during the initiation stage in *A. thaliana* and cotton (Bui *et al*., 2019). *GLABROUS1* (*GL1*, a MYB transcription factor [TF] gene), *GLABROUS3* (*GL3*, a bHLH TF gene), *ENHANCER OF GLABRA3* (*EGL3*, a bHLH TF gene) and *TRANSPARENT TESTA GLABRA1* (*TTG1*, a WD40 TF gene) have been suggested to play regulatory roles in the trichome development through the induction of *GLABRA2* (*GL2*, a HD-ZIP IV TF gene)(Pesch *et al*., 2015). In addition, *GhMYB25*, another type of *MYB* gene, was found to play a major role in cotton fiber initiation and extension, and similar functions were found for its homologues, *GhMYB25-LIKE, GhMML3_A12* and *GhMML4_D12* (Wu *et al*., 2018). *GbML1*, another HD-ZIP IV TF, was found to have a similar expression pattern to *GhMYB25* and to interact with *GhMYB25* to control fiber development (Zhang *et al*., 2010). As a model of rapid single-cell growth, sucrose metabolism-related proteins also play important roles in the process (Ruan, 2014).

Sucrose is an important form of transportable sugar in plants. Plants convert CO_2_ into organic carbon through photosynthesis in leaves, and then transport it to heterotrophic sinks, such as the stem, root and fruit, for energy storage and utilization (Ruan, 2014). Thus, sucrose metabolism plays a critical role in plant growth and development, including trichome development. Sucrose can be synthesized by sucrose phosphate synthase and sucrose phosphate phosphatase, and degraded by sucrose synthase or invertase in plants (Ruan, 2014). Sugar signals produced by invertase likely influence the cotton fiber initiation through MYB TFs and auxin signals (Wang *et al*., 2014). High turgor pressure induced by sucrose degradation via invertase and sucrose synthase promotes the rapid elongation of cotton fibers (Wang *et al*., 2010, Ahmed *et al*., 2018). In contrast to invertase, uridine diphosphate (UDP)-glucose produced by sucrose synthase is a vital substrate for cellulose synthesis, and it influences cell wall synthesis, especially of the secondary cell wall (Ahmed *et al*., 2018, Fujii *et al*., 2010). Additionally, two overexpression studies showed that sucrose phosphate synthase has an effect on fiber development (Park *et al*., 2008, Haigler *et al*., 2007),which presumably because sucrose phosphate synthase can recycle fructose released by sucrose synthase to obtain more sucrose (Martin and Haigler, 2004). Similar to cotton fibers and *Arabidopsis* epidermal hairs, in essence, poplar catkin fibers also belong to plant trichomes. However, compared with the numerous important advances in *Arabidopsis* and cotton, the research foundation of catkin fibers development in poplar is quite weak. Whether sucrose metabolism has a close relationship with the poplar catkin fibers development and what are the real roles of candidate genes in poplar, both of which require further exploration.

Chinese white poplar (*Populus tomentosa* Carr.) is a representative species of poplar, widely planted in the eastern region of Asia. We previously conducted transcriptome analysis of catkin fibers development in this plant, and found that many Gene Ontology (GO) terms related to sucrose metabolism were significantly enriched during this process (Ye *et al*., 2014). Therefore, in this study, we made histological observations of catkin fibers development in *P. tomentosa*, and measured the sugar content and activities of key sucrose metabolism enzymes at various developmental stages. Using the genomic data of *P. tomentosa* compiled by our group (An *et al*., 2022), we searched for sucrose metabolism-related genes and performed detailed fluorescence-based real-time quantitative polymerase chain reaction (RT-qPCR) analysis. Additionally, the expression patterns of key potential candidate genes involved in poplar catkin fibers development were analyzed, and a gene co-expression network was constructed. The relationships between sucrose metabolism-related genes and TFs were also analyzed. Finally, two key genes, *PtoSUS2* and *PtoVIN3*, were selected to investigate the function of sucrose metabolism in catkin fibers formation using the model plant *A. thaliana*.

## Materials and methods

### Plant materials and histological analysis

Live branches of adult female *P. tomentosa* were collected in February 2018, at which time the flower buds were undergoing megasporogenesis and embryo sac formation (An, 2010). The branches were placed in a greenhouse (25 ± 2°C, 70% humidity) for hydroponic culture, and samples were collected as described previously (Ye *et al*., 2014). The capsules which stigmas wilted and no longer secreted mucus at 2 days post-anthesis (DPA) were collected and represented as 0 h, and then the capsules were collected when they were continuously cultured for 24, 34, 48, 58, 72, 96 and 120 h. Samples collected at these eight time points were subjected to histological analysis; according to the analysis results, the samples from the last five time points, which were related to catkin fibers development, were subjected to RNA extraction as well as sugar content and enzyme activity analyses. RNA extraction and histological analysis were also performed as described previously (Ye *et al*., 2014).

The Columbia ecotype (Col) and two trichome mutants, *trichome less 1* (*tl1*, SALK_133117C) and *trichome less 2* (*tl2*, SALK_039825C), were used in this study. The seeds were sterilized in 5% sodium hypochlorite solution for 15 min, and then placed in half-strength Murashige-Skoog (MS) medium at 4 [for 72 h. Subsequently, the seeds were cultivated in an incubator at 22 [under LD (16 h light/8 h dark) condition for 7 days, then transplanted into nutrient soil with added perlite.

### Quantitation of sugar content and enzyme activities

Extraction of soluble sugars is described in detail in the Methods S1. The concentrations of sucrose, fructose and glucose were analyzed by high-performance liquid chromatography (HPLC) using an Agilent 1260 HPLC system (Agilent Technologies, Santa Clara, CA, USA), and a carbohydrate column (4.6 × 150 mm, 5 μm, Agilent Technologies). The sugar concentrations were quantified according to a standard solution (Sigma, St. Louis, MO, USA), and defined in terms of mg•g^-1^ DW.

The activities of sucrose synthase and sucrose phosphate synthase were analyzed according to the method of Yang et al. (Yang *et al*., 2013) with some modifications (described in detail in Methods S1). The activities of neutral/alkaline, vacuolar, and cell wall invertases were analyzed according to previously described methods (Chen *et al*., 2015). The enzyme activities were recorded as μmol•g^-1^ FW•h^-1^.

### Candidate gene identification and RT-qPCR

To determine the expression levels of key genes during the development of catkin fibers, primers were designed to amplify fragments specific to each gene. Genes from *A. thaliana* and cotton (Table S1) were used as queries for a Basic Local Alignment Search Tool for Proteins (BLASTP) search against the protein sequences from the *P. tomentosa* genomic data (An *et al*., 2022). The sequences of the primer pairs used for each gene are listed in Table S2.

Total RNA was used for first-strand cDNA synthesis with the Reverse Transcription System I (Promega, Madison, WI, USA), then diluted 1:50 in water. The diluted cDNA was used as a template, and PCR amplification was performed with the 7500 Fast Real-Time PCR System (Applied Biosystems, Foster City, CA, USA) using the following program: 94 for 5 min (one cycle); 94 for 30 s, 60 for 20 s, 72 for 30 s (35 cycles); 72 for 5 min. The *PtActin* gene was used as an internal reference gene, as described previously (Chen *et al*., 2018), and the SYBR Premix Ex Taq kit (Takara, Shiga, Japan) was used as the reaction solution. All analyses were performed with three technical and three biological replicates.

### Phylogenetic tree and co-expression network

To confirm the evolutionary relationships in sucrose metabolism related-gene families, all genes were compared with their homologues from poplar*, A. thaliana* and cotton through multiple sequence alignment (Table S3) using the ClustalW program within MEGA 7.0 software. Phylogenetic trees were draw with MEGA 7.0 using the neighbor-joining method with complete deletion, and 1,000 replications were used for bootstrap analysis. Gene co-expression network analysis was performed to identify correlated genes using R software (v3.6.3), and the networks were visualized using Cytoscape (v.3.7.1).

### Yeast one-hybrid assay (Y1H)

To characterize the interaction between *PtoSUS2*/*PtoVIN3* and potential TFs, the promoter region of *PtoSUS2* and *PtoVIN3* were amplified and cloned into the vector pAbAi-*proPtoSUS2* and pAbAi-*proPtoVIN3*. Thereinto, according to the results based on PlantCARE and New PLACE, the promoter region containing *MYB* and *bHLH* motifs were cloned and divided into two regions, and the potential TFs (Table S4) were amplified and cloned into the vector pGADT7. Subsequently, the two vectors were successively transferred into the Y1H Gold yeast strain according to the Y1H transformation protocol (Shanghai Weidi Biotechnology Co., Ltd). Primers used for Y1H were shown in Table S5.

### Transformation and gene expression analysis in *A. thaliana*

The open reading frame sequences of *PtoSUS2* and *PtoVIN3* were amplified and cloned into the binary vector pCAMBIA 1301 to generate overexpression vectors. Primers used for over-expression vector construction were shown in Table S5. Transgenic lines were obtained by the *Aarobacterium tumefaciens*-mediated method established by (Zhang *et al*., 2006). Transgenic plants were selected on half-strength MS medium containing 25 mg/L hygromycin. Gene expression analysis was performed as above, except that relative expression levels were calculated using the 2^-ΔΔCt^ method. The sequences of the primer pairs used for trichome-related genes in *A. thaliana* and exogenous genes are listed in Table S2.

### Morphological analysis and scanning electron microscopy (SEM)

The sixth rosette leaves, stems (morphological bottom), and bottom-most cauline leaves were used for trichome comparison with a stereomicroscope (Stemi 305, ZEISS). The number and length of trichomes in rosette leaves was determined in plants that had been cultured for 30 days after germination (DAG), and the trichome length were analyzed according the previous study (Li *et al*., 2013b), and the primary root length and lateral root number were quantified at 10 DAG. Five biological replicates were performed. For SEM, the samples were cut into 1 × 1-cm pieces, and dehydrated in a vacuum freeze dryer (FreeZone 4.5L, Labconco) for 48 h. Images were obtained using a scanning electron microscope (S-3400N, Hitachi, Japan).

### Statistical Analysis

Sugar content, enzyme activity and gene expression data were analyzed through one-way analysis of variance (ANOVA) using SPSS software (version 22.0 for Windows; SPSS Inc., Chicago, IL, USA). The correlations between genes expression patterns in sucrose metabolism and sugar contents was analyzed by Pearson correlation using the same software. Two-tail test was used to test for significance level as *P* < 0.05 and *P* < 0.01.

### Accession numbers

The gene sequences can be found under the accession numbers: *PtoSUS2* (ON534005), *PtoVIN3* (ON534006).

## Results

### Morphological characteristics of poplar catkin fibers

The morphology of catkin fibers changed rapidly during hydroponic culture (Fig. 2). Capsules obtained at the first time had two ovaries with ovule inverted on placenta and no funicle could be seen (Fig. 2a). Histological analysis revealed that the size of ovule gradually increases, the funicle elongated and expanded, and the placenta expanded but no catkin fibers appeared when the branches had been cultured for 0-48 h (Fig. 2a-d). The epidermal cells of funicles initially showed larger nuclei (Fig. 2e, f), in accordance with the endoreduplication stage in *A. thaliana*, indicating the initial stage of development of catkin fibers. Subsequently, the epidermal cells exhibited protrusion and extension from 58 to 120 h, until the catkin fibers finally full filled the ovary, while there was no obvious morphological difference in ovule, funicle and placenta (Fig. 2g-j). Thus, according to the morphological characteristics of catkin fibers, we divided the development process into five stages: stage 1 (preparatory stage, cultured for 48 h, Fig. 2d), stage 2 (initial stage, cultured for 58 h, Fig. 2e-f), stage 3 (apophysis stage, cultured for 72 h, Fig. 2g-h), stage 4 (extension stage, cultured for 96 h, Fig. 2i), stage 5 (maturation stage, cultured for 120 h, Fig. 2j). Furthermore, as shown in Fig. 2, catkin fibers in poplar were produced from the epidermal cells of funicle and placenta, independent from seeds, which was totally different from cotton fibers. Samples from the five stages were used for analyses of enzyme activities and gene expression patterns.

**Figure 2.**
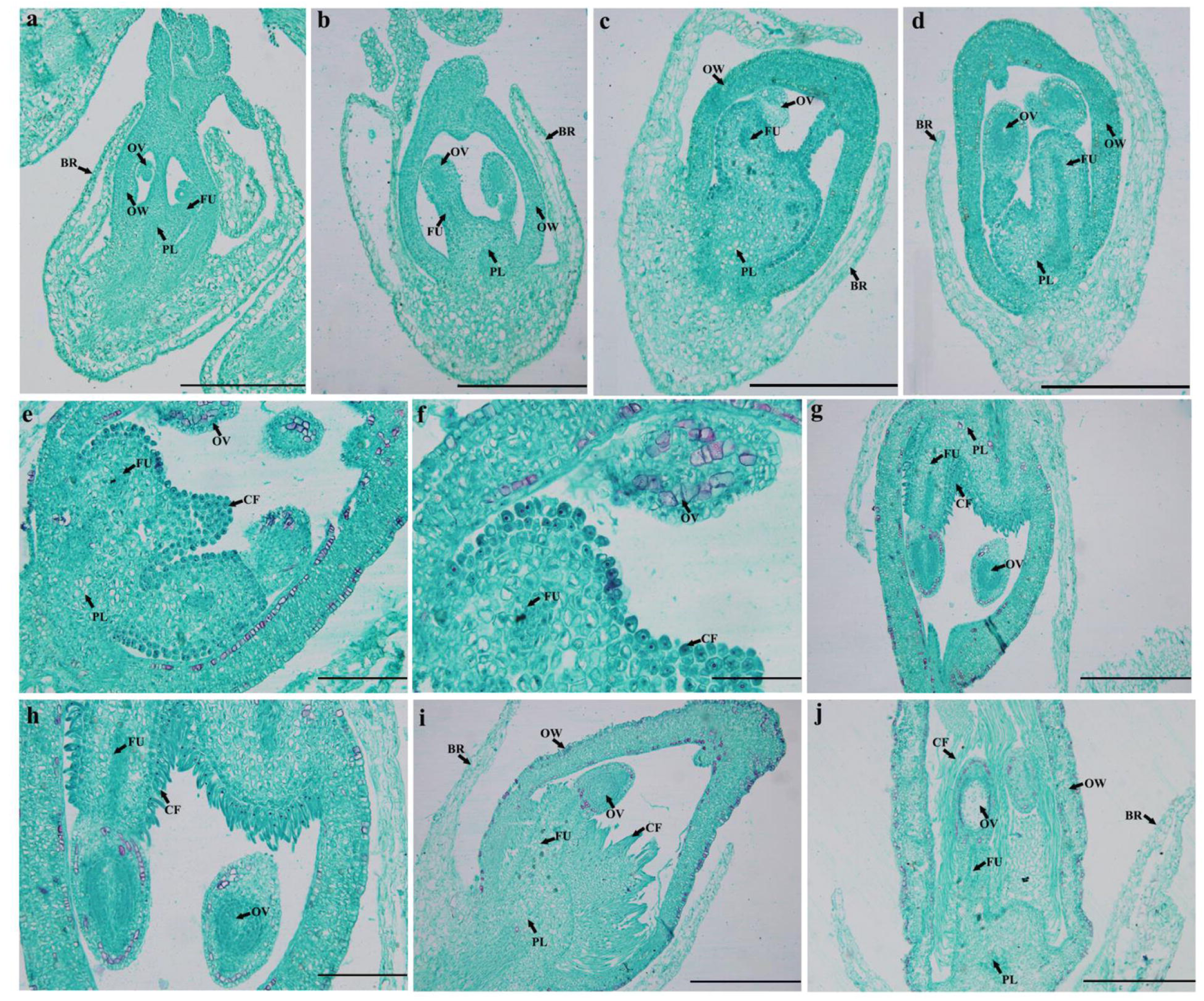
Histological characteristics of female buds in *P. tomentosa.* a-d. Female buds on branches that were hydroponically cultured for 0, 24, 34, 48 h, respectively. No catkin fibers were observed. **e, f** Female buds on branches that were hydroponically cultures for 58 h. **g, h** Female buds on branches that were hydroponically cultured for 72 h. **i** Female buds on branches that were hydroponically cultured for 96 h. **j** Female buds on branches that were hydroponically cultured for 120 h. OW, ovary wall, OV, ovule, FU, funicle, BR, bract, CF, catkin fibers. PL, placenta. Scale bar: **a-d, g, i, j** = 500 μm, **e, h** = 200 μm, **f** = 100 μm.

### Changes in sugar content and key enzyme activities in poplar catkin fibers

The sucrose content of the catkin of *P. tomentosa* showed a downward trend, indicating that sucrose is gradually consumed during the development of catkin (Fig. 3a). In particular, during the stage 3-4, sucrose was consumed at a dramatic rate when the catkin fibers extended rapidly. Glucose and fructose showed similar trends, which both had higher contents during stage 2-4. The ratio of hexose to sucrose was also analyzed. As shown in Fig. 3b, the ratio increased sharply between the apophysis stage (stage 3) and extension stage (stage 4), and decreased between the extension stage (stage 4) and maturation stage (stage 5). Changes in the ratio indicated that the high hexose concentration may play a role in catkin fibers extension. When the catkin fibers began to extend, most of the sucrose present was degraded into glucose and fructose, which likely provided greater turgor pressure, thus promoting catkin fibers elongation. To further explore the role of sucrose in catkin fibers development, we also determined the activities of key enzymes in sucrose metabolism. Consistent with the results of sugar content analysis, sucrose degradation was the dominant process.

**Figure 3.**
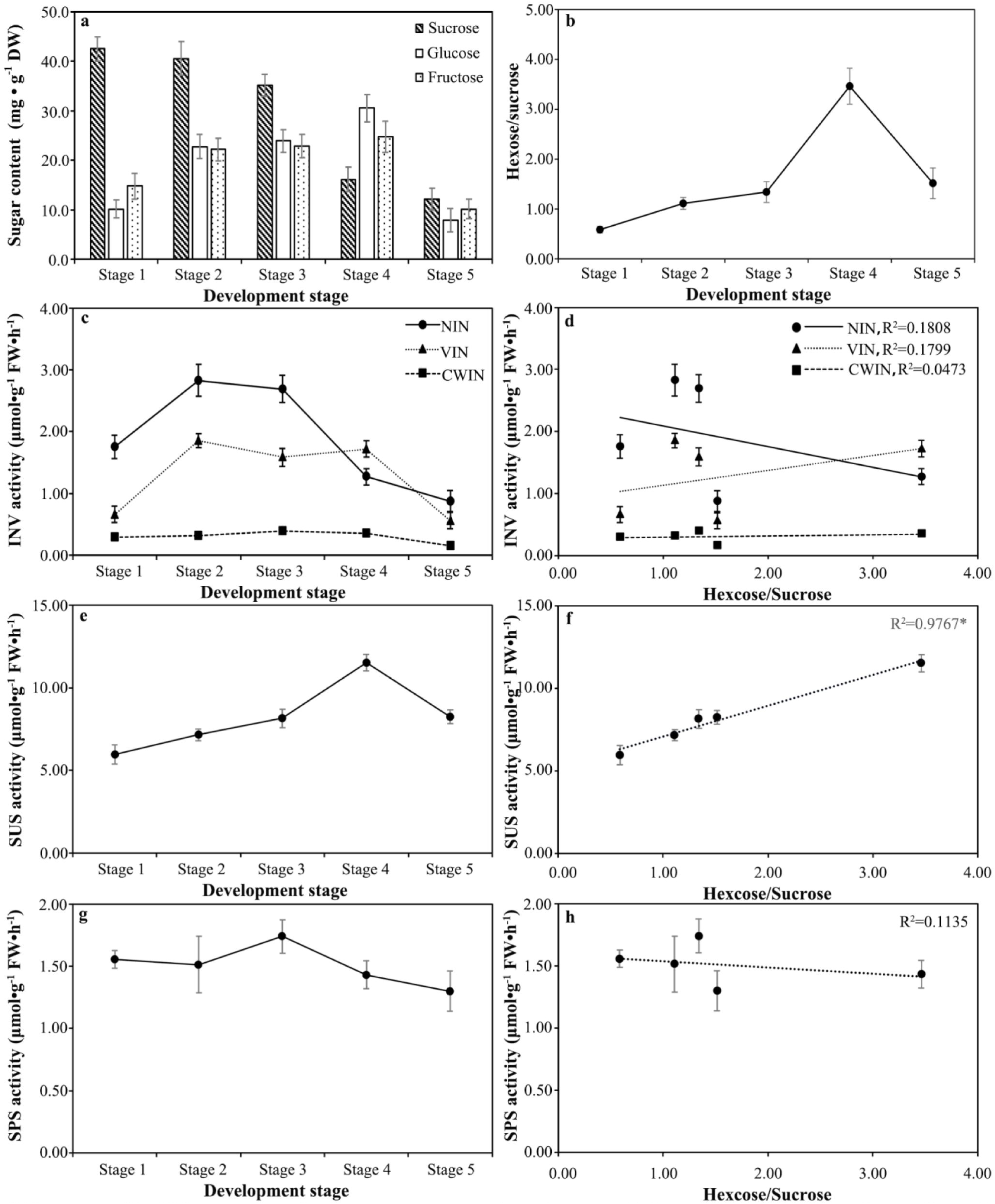
Changes in sugar contents and key enzymes involved in sucrose metabolism during poplar catkin fibers development. Bars represent the standard deviation (SD, n = 3).

The activity of cell wall invertase showed no significant fluctuations except at stage 5, when it decreased as the catkin fibers matured (Fig. 3c). The activities of neutral/alkaline invertase and vacuolar invertase were higher than that of cell wall invertase. Vacuolar invertase activity increased during catkin fibers initiation and decreased during the catkin fibers maturation, indicating that vacuolar invertase may play a role in catkin fibers development (Fig. 3c). Meanwhile, the activity of neutral/alkaline invertase was high only during the initial (stage 2) and apophysis stages (stage 3), and appeared to play a role in the early development of catkin fibers (Fig. 3c). Compared with invertase, sucrose synthase showed higher activity during the development of catkin fibers, up to the extension stage (stage 4); it then decreased during the maturation stage (stage 5), showing an inverse correlation with sucrose content (Fig. 3e). Similar results were obtained though fitting analysis. Sucrose synthase activity showed a significant correlation with the ratio of hexose to sucrose, indicating that sucrose synthase plays a major role in sucrose degradation during the development of catkin fibers (Fig. 3f). Furthermore, sucrose phosphate synthase activity was analyzed and used to represent the sucrose synthesis pathway. As shown in Fig. 3g, sucrose phosphate synthase exhibited steady activity throughout catkin fibers development, except during the apophysis stage (stage 3).

### Gene expression patterns related to sucrose metabolism in poplar catkin fibers

The expression of *cell wall invertase* (*CWIN*) showed a similar trend to enzyme activity. We analyzed three members of the gene family, *PtoCWIN3*, *PtoCWIN4* and *PtoCWIN5*, during catkin fibers development. *PtoCWIN3* appeared to play a major role in catkin fibers development based on its high and variable expression level (Fig. 4a). However, Pearson correlation analysis showed that *PtoCWIN4* rather than *PtoCWIN3* was positively correlated with sucrose content, which may be due to the dramatic decreased in *PtoCWIN3* at the initial stage (Table 1). Two members of the *VIN* gene family were observed, and both had elevated expression at stage 2-4, coinciding with the enzyme activity results (Fig. 4b). And both members of the *VIN* gene family had positive correlations with the concentrations of fructose and glucose (Table 1). For *neutral/alkaline invertase* (*NIN*), nine members were detected during catkin fibers development, including two low-expression members, *PtoNIN2* and *PtoNIN11* (Fig. 4d, f). According to their expression patterns, the other members can be divided into two groups. *PtoNIN3* and *PtoNIN4* had higher expression at the apophysis stage (Fig. 4e), while the other five members peaked at the initial stage (Fig. 4c-d, f-g). It is worth noting that the expression patterns of members with close evolutionary relationships are not completely consistent (Fig. S1).

**Figure 4.**
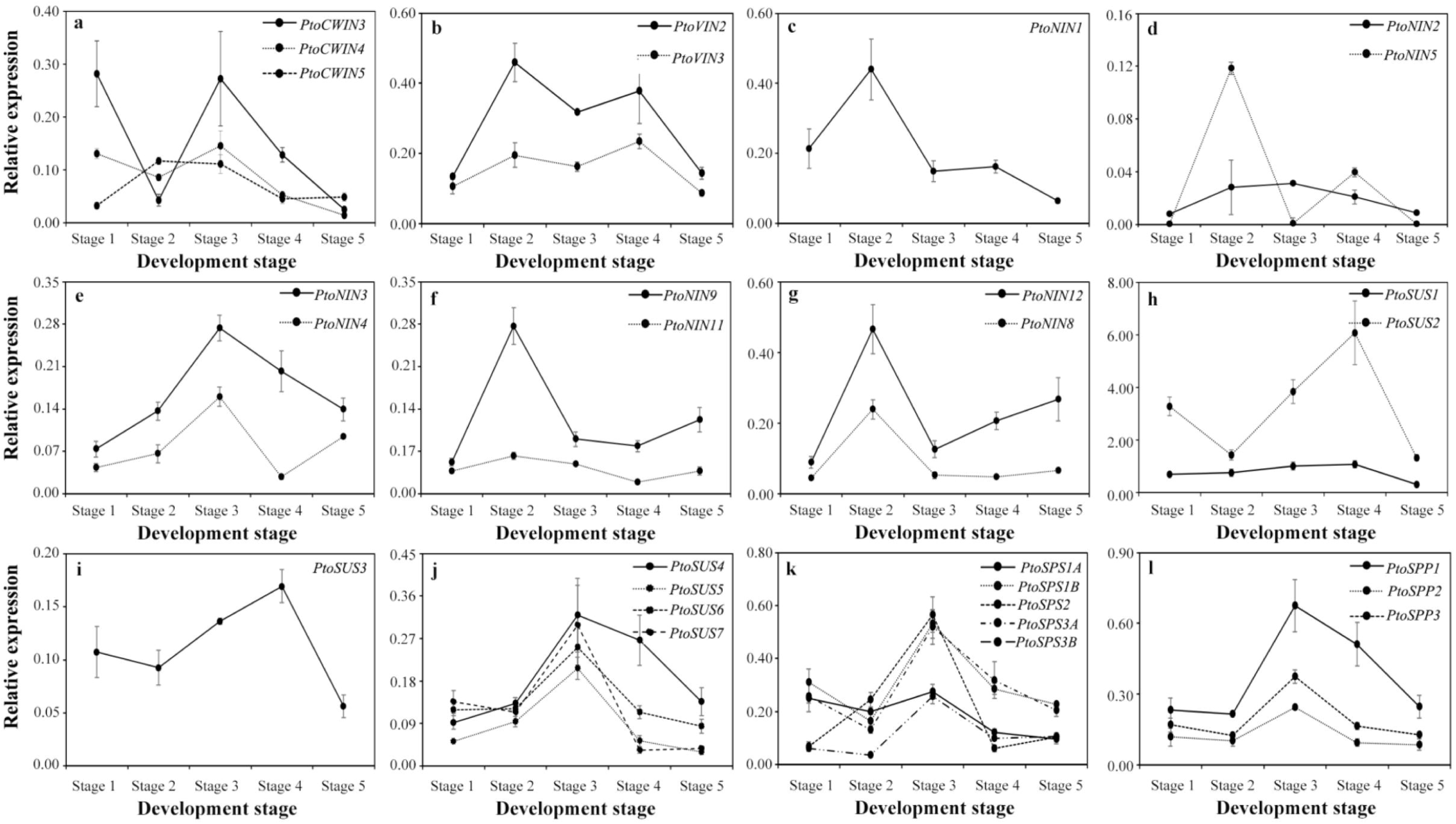
Analysis of sucrose metabolism related gene families in the context of poplar catkin fibers development. **a** Expression analysis obtained by RT-qPCR for *CWIN* gene family. **b** Expression analysis obtained by RT-qPCR for *VIN* gene family **c-g** Expression analysis obtained by RT-qPCR for the *NIN* gene family. **h-j** Expression analysis obtained by RT-qPCR for the *SUS* gene family. **k, l** Expression analysis obtained by RT-qPCR for *SPS* and *SPP* gene family, respectively. Bars represent the ± SD.

**Table 1.**
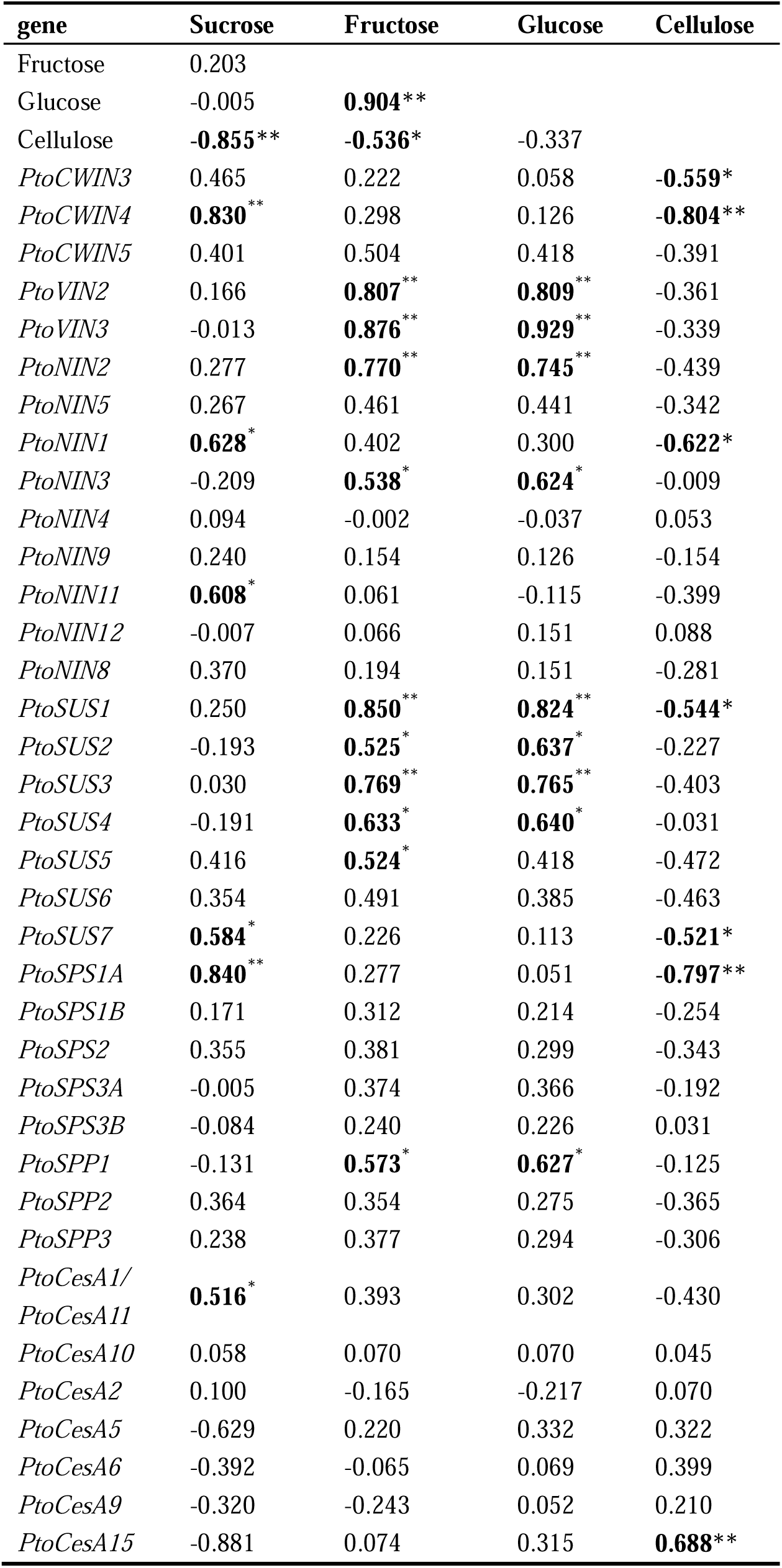
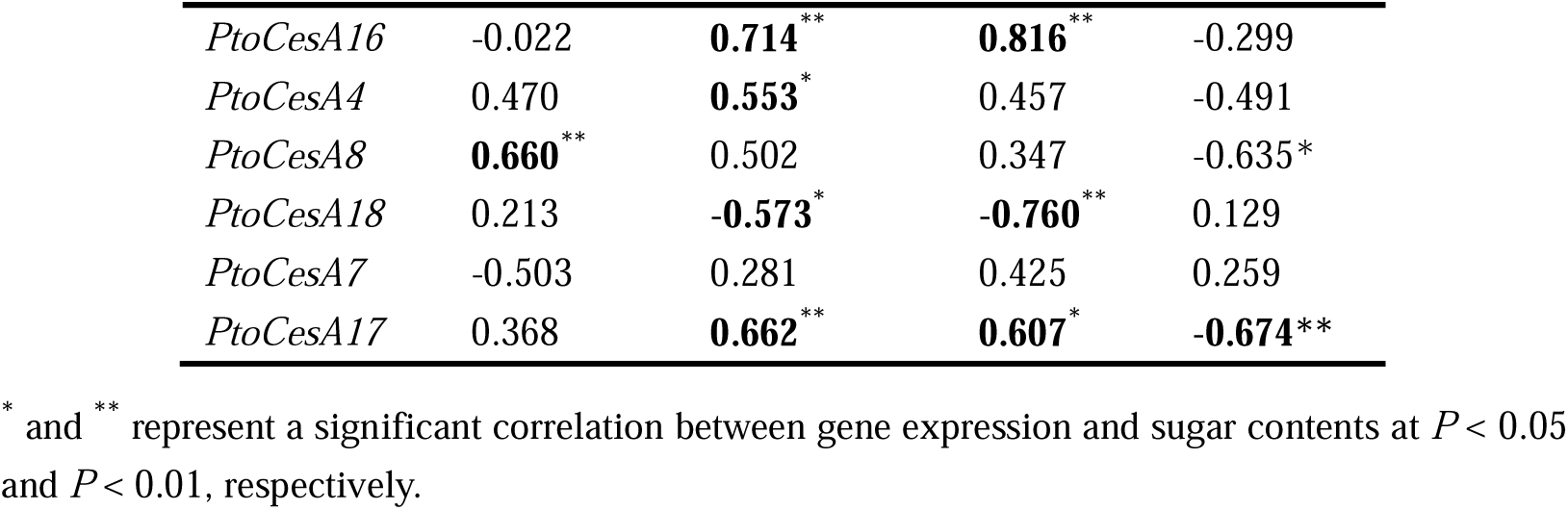
Correlations between the expression patterns of genes involved in sucrose metabolism and sugar contents in poplar catkin fibers development.

There are seven members in the *SUS* gene family, all of which were detected during poplar catkin fibers development. Their expression pattern showed a clear correlation with the phylogenetic tree (Fig. S1c). The first cluster, comprising *PtoSUS1* and *PtoSUS2*, had high expression during catkin fibers development. Compared with *PtoSUS1*, *PtoSUS2* had higher expression, and both had their highest levels during the extension stage (Fig. 4h). The second cluster only contains one member, *PtoSUS3*, which had low expression but a similar pattern to the first cluster (Fig. 4i). The other four members belonged to the third cluster, and had high expression at the apophysis stage (Fig. 4j). The expression levels of most *SUS* gene members had significant positive correlations with the concentration of hexose (Table 1).

The expression patterns of sucrose phosphate synthase and sucrose phosphate phosphatase gene families were analyzed. Five of the six members of the *SPS* gene family were analyzed, and they all had high expression at the apophysis stage, in accordance with the results for sucrose phosphate synthase activity (Fig. 3g and 4k). All three sucrose phosphate phosphatase members (*PtoSPP1*, *PtoSPP2* and *PtoSPP3*) were detected, and all had elevated expression during the apophysis stage (Fig. 4l). The results of both the enzyme activity and gene expression analyses showed that the sucrose synthesis pathway might only have a slight effect at the apophysis stage.

### *CesA* gene family and other potential genes in poplar catkin fibers development

Poplar catkin fibers follows a similar development process to cotton fiber, and cellulose is the main component of cotton fiber (Fujii *et al*., 2010). Thus, we also checked the expression pattern of the *cellulose synthase* (*CesA*) gene family in poplar catkin fibers. Thirteen members of the *CesA* gene family were analyzed in this study, which belonged to five of six groups on the phylogenetic tree, i.e., all except cluster P2 (Fig. S1d). Among them, *PtoCesA17*, belonging to clade P1, had high expression at the apophysis stage, while its homologue *PtoCesA7* had low expression (Fig. 5d). Homologous pairs in clade P3 had similar expression patterns, except *PtoCesA2* and *PtoCesA5* (Fig. 5e-g). *PtoCesA6* and *PtoCesA9* had high expression throughout the process of catkin fibers growth (Fig. 5g). *PtoCesA15* appeared to play a more important role than *PtoCesA16*, as it had dramatically higher expression during catkin fibers development (Fig. 5f). Other genes belonging to clades S1, S2 and S3, had low expression in catkin fibers, although their expression increased at the initial and apophysis stages with the exception of *PtoCesA18* (Fig. 5a-c). Notably, the cellulose content in poplar catkin fibers gradually increased as the development process (Methods S2; Fig. S2). Correlation analysis showed that cellulose content has a negative relationship with sucrose content, and positive relationship with the expression of *PtoCesA15* (Table 1).

**Figure 5.**
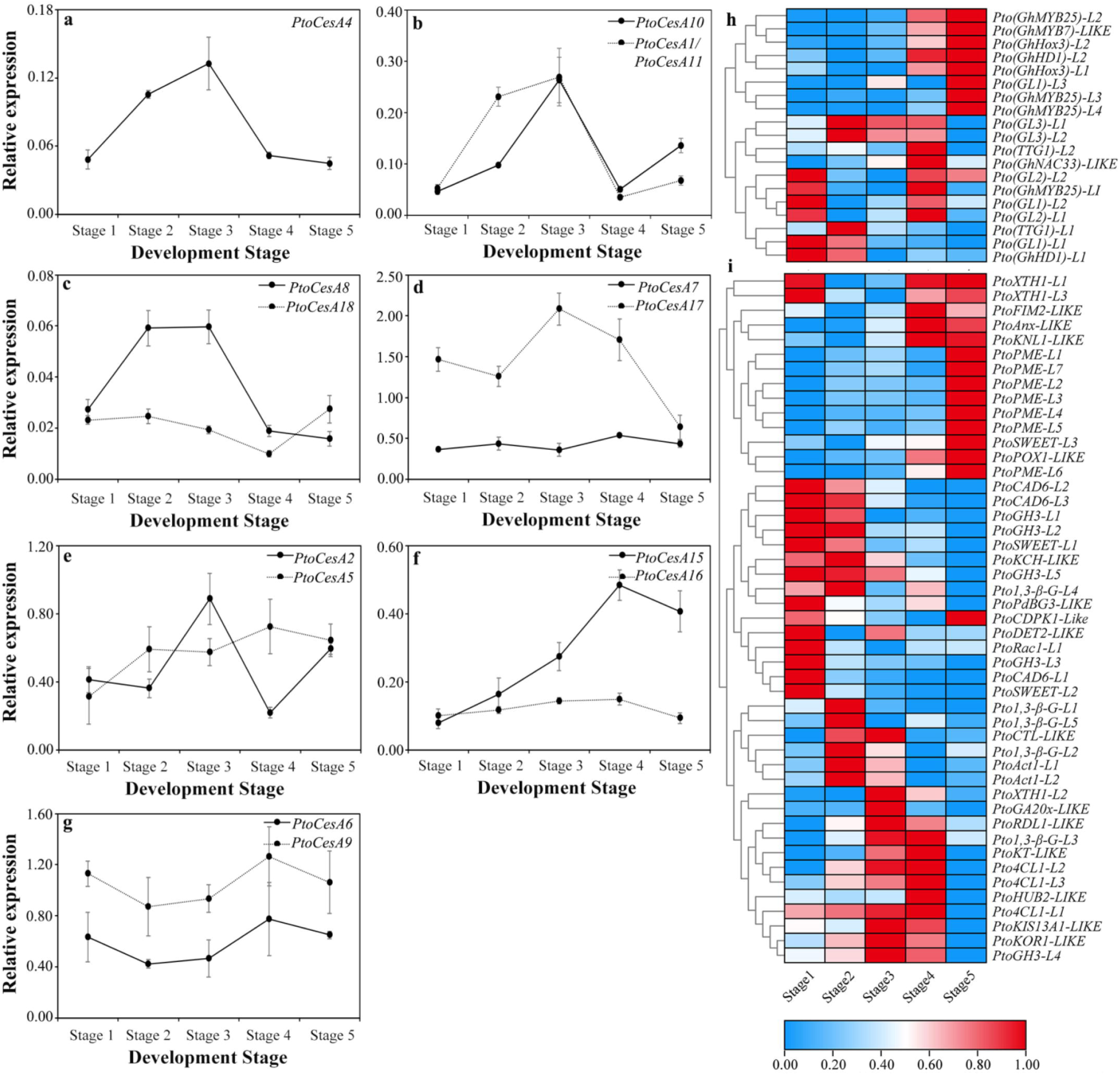
Analysis of the *CesA* gene family and other potential genes in the context of poplar catkin fibers development. **a-g** Expression analysis based on RT-qPCR for *CesA* gene family. **h, i** Heat map of gene expression obtained by RT-qPCR for TFs-related and other genes, respectively. Bars represent the ± SD.

The expression patterns of potential TFs (Table S1) were also analyzed. Thereinto, these homologues can be divided into three subgroups (Fig. 5h). Among bHLH TFs, *Pto(GL3)-L1* and *2* had highest expression at the initiation stage. In the *WD40* gene family, *Pto(TTG1)-L1* had high expression at the initiation stage, while the another WD40 TF *Pto(TTG1)-L2* had high expression at the extension stage. Members of the MYB gene family may have divergent functions during catkin fibers development. *Pto(GL1)-L1* and *2* had high expression during early development of catkin fibers, and *Pto(GL1)-L2* had an additional role in the extension stage. In contrast, *Pto(GL1)-L3* showed increased expression at the maturation stage. The homologues of *GhMYB25* all had increased expression at the maturation stage, except *Pto(GhMYB25)-L1*. *Pto(GhMYB7)-LIKE*, the cotton homologue of which was speculated to be a lipid transfer protein-related gene, also had high expression at the maturation stage. Another important TF gene family, HD-ZIP IV, was also analyzed during poplar catkin fibers development. *Pto(GL2)-L1* and *2* had high expression at the preparatory and extension stages, and the homologues of *GhHox3* had functions at the maturation stage. Intriguingly, opposing results were found between *Pto(GhHD1)-L1* and *Pto(GhHD1)-L2*.

We also examined the expression of other potential key candidate genes in poplar (Table S1). As shown in Fig. 5i, three subgroups played different roles in the early, middle and late development stages. In cluster I, *PtoPME-related* genes played an important role in catkin fibers maturation, which indicted dramatic changes in the composition of cell wall. In cluster II, *PtoCAD6-related* and *PtoGH3-related* appeared to be important to the early development of catkin fibers. During the middle development stage (cluster III), many genes were involved. Thereinto, *PtoAct1-L1* and *2* and *Pto1,3-β-G-L1* and *5* had high expression at the initial stage. *PtoGA20x-LIKE, PtoKIS13A1-LIKE, PtoKOR1-LIKE* and *PtoXTH1-L2* functioned mainly at the apophysis stage, while *PtoHUB2-LIKE, Pto1,3-β-G-L3, Pto4CL1-L2, Pto4CL1-L3* and *PtoKT-LIKE* had high expression at the extension stage.

### Co-expression network of genes involved in poplar catkin fibers development

To further explore the relationship between sucrose metabolism and poplar catkin fibers development, co-expression network analysis was performed using sucrose metabolism-related genes and potential key genes involved in poplar catkin fibers development. As shown in Fig. 6a, there were four divergent gene clusters, and sucrose metabolism-related genes mainly occurred in cluster I-III. Thereinto, cluster II was mainly composed of sucrose synthesis-related genes, with *PtoSPS2* playing a leading role. Notably, *PtoSUS5-7*, which is a homologue according to the phylogenetic tree, and *PtoGA20x-LIKE* had close relationships with *PtoSPS* and *PtoSPP*. Different members of the *CesA* gene family play specific roles in clusters I and III, respectively. Cluster IV represents the group of genes that function in the late stages of catkin fibers development. Additionally, we performed network analysis between sucrose metabolism-related genes and TFs using data from a previous study (Ye *et al*., 2014). As shown in Fig. 6b, many TF gene families were involved in poplar catkin fibers development, including *bHLH*, *ERF*, *WRKY*, *MYB* and *HD-ZIP*. *Cis*-acting elements were also predicted for genes related to sucrose metabolism in this study (Fig. S3). Thereinto, MYB and MYC motifs were found to be widely distributed in the promoter region of *PtoSUS2* and *PtoVIN3* (Fig. 6c). Therefore, the MYB and bHLH TFs (Table S5), which had strong correlation with the expression of *PtoSUS2* and *PtoVIN3*, were selected to do Y1H assay (Fig. 6d). For *Pro-PtoSUS2-2*, strong interactions were found for some MYB TFs, including *Pto(GL1)-L3*, *Pto(MYB25)-L2/4* and *Pto(DRMY1)-LIKE*, and strong interactions were found between *Pro-PtoSUS2-1* and *Pto(GhMYB25)-L4*. However, only slightly interactions were found in *Pro-PtoVIN3-2*, both with MYB and bHLH TFs.

**Figure 6.**
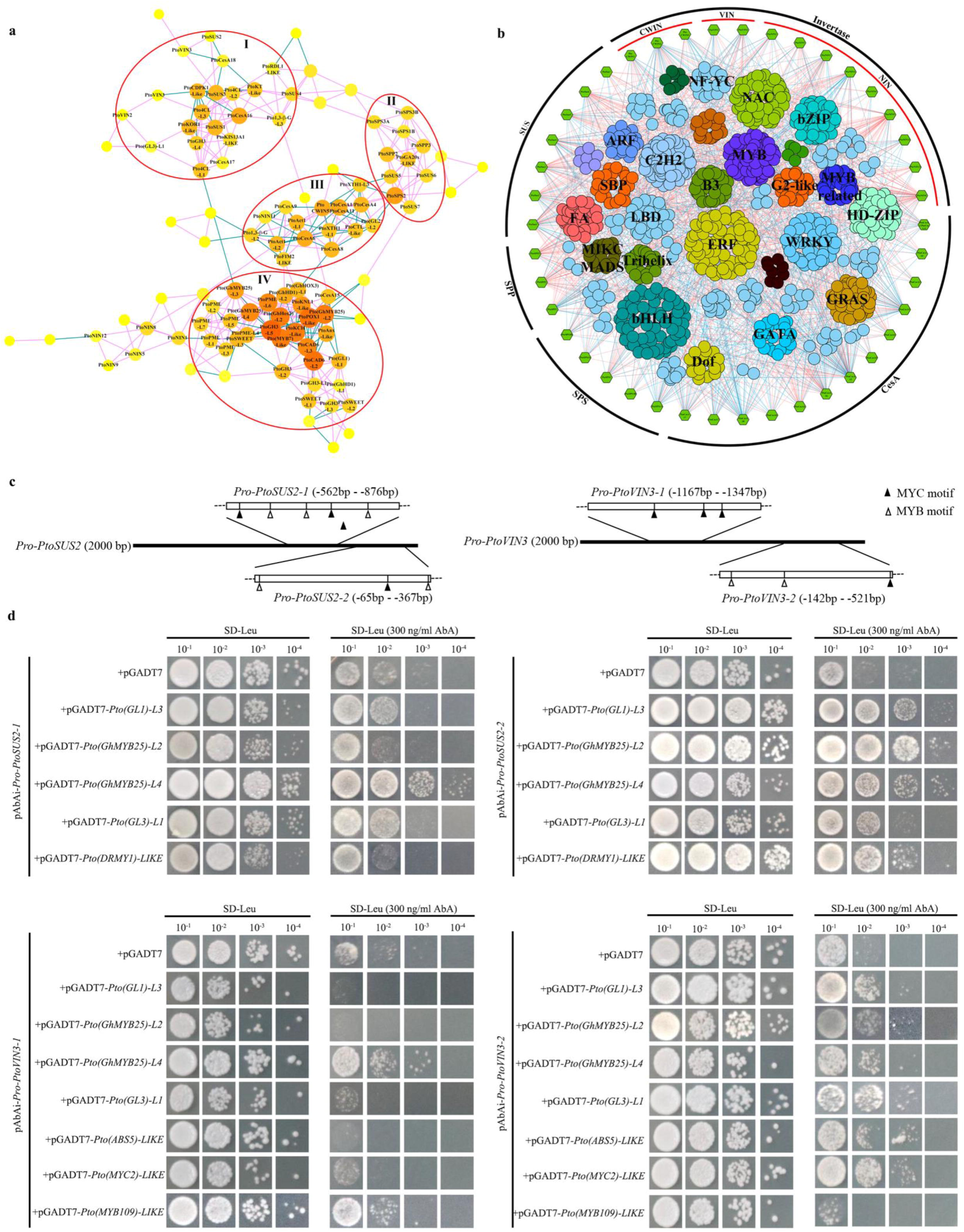
Co-expression network analysis. **a** Co-expression network of 108 candidate genes associated with catkin fibers development in *P. tomentosa*. The threshold for significance of the Pearson correlation coefficient was set to ≥ 0.8 (or ≤ − 0.8), p-value ≤ 0.05. Larger and redder the circles, indicate higher connectivity. **b** Co-expression network of sucrose metabolism-related genes and TFs, the threshold for significance of the Pearson correlation coefficient was set to ≥ 0.8 (or ≤ − 0.8), p-value ≤ 0.01. **c** MYB and MYC motifs in the promoter regions of *PtoSUS2* and *PtoVIN3*. **d** Activity analysis of the yeast transformed with the promoter of *PtoSUS2*/*PtoVIN3* and TFs.

### Overexpression of *PtoSUS2* and *PtoVIN3* increases trichome density in wild-type *A. thaliana* and *tl1/2* mutant

The results for sucrose synthase and vacuolar invertase indicate roles for sucrose metabolism in poplar catkin fibers development; two key genes, *PtoSUS2* and *PtoVIN3*, were selected to demonstrate the exact role. Thereinto, *PtoSUS2* had a stable expression level except in root, while *PtoVIN3* had higher expression levels in stem and root, and the two genes all played roles during the early development of female and male floral bud (Fig. S4). Because of the feasibility and long period involved in generating transgenic poplars and investigating the phenotype of poplar catkins, the *A. thaliana* trichome system was used to analyze gene function. We introduced *PtoSUS2* and *PtoVIN3* constructs driven by the 35S promoter into wild-type *A. thaliana* and used T3-generation plants to analyze the phenotype. Thirteen and twenty-two independent transgenic lines were finally obtained for *PtoSUS2* and *PtoVIN3*, respectively. Three lines of each transgenic type were extensively evaluated. In wild-type *A. thaliana*, trichomes are widely distributed on the adaxial and abaxial surfaces of rosette leaves and cauline leaves, as well as stems. Compared with the wild-type, the *PtoSUS2* and *PtoVIN3* overexpression lines had higher leaf trichome densities on the adaxial surface of rosette leaves and stems at 30 DAG, and trichome shape was unaffected (Fig. 7a-f; Fig. S5). The number of sixth-rosette leaf trichome in the wild-type was 67 ± 8.80, significantly fewer than in the transgenic plants, *Pro35S:PtoSUS2/col-2* (87 ± 6.89) and *Pro35S:PtoVIN3/col-24* (94 ± 8.41) (Fig. 7g), and ectopic expression of *PtoSUS2* and *PtoVIN3* in *Arabidopsis* also significantly increased the leaf trichome length (Fig. 7h). However, the rosette leaf number, fresh weight and height did not significantly differ between wild-type and overexpression plants (Fig. S6g-i). Similar results were obtained in terms of rosette leaf shape (20 DAG) and whole plants (40 DAG) (Fig. S6a-f), while the primary root length and lateral root number (10 DAG) were slightly increased in overexpression plants, especially in *Pro35S:PtoSUS2/col-2* (Fig. 7i, j; Fig. S6). Additionally, the trichome density of transgenic plants on the abaxial surface of rosette leaves slightly increased, but the numbers of trichomes on the adaxial and abaxial surfaces of cauline leaves had no significant difference, and these results were not affected by incubation time (Fig. S7).

**Figure 7.**
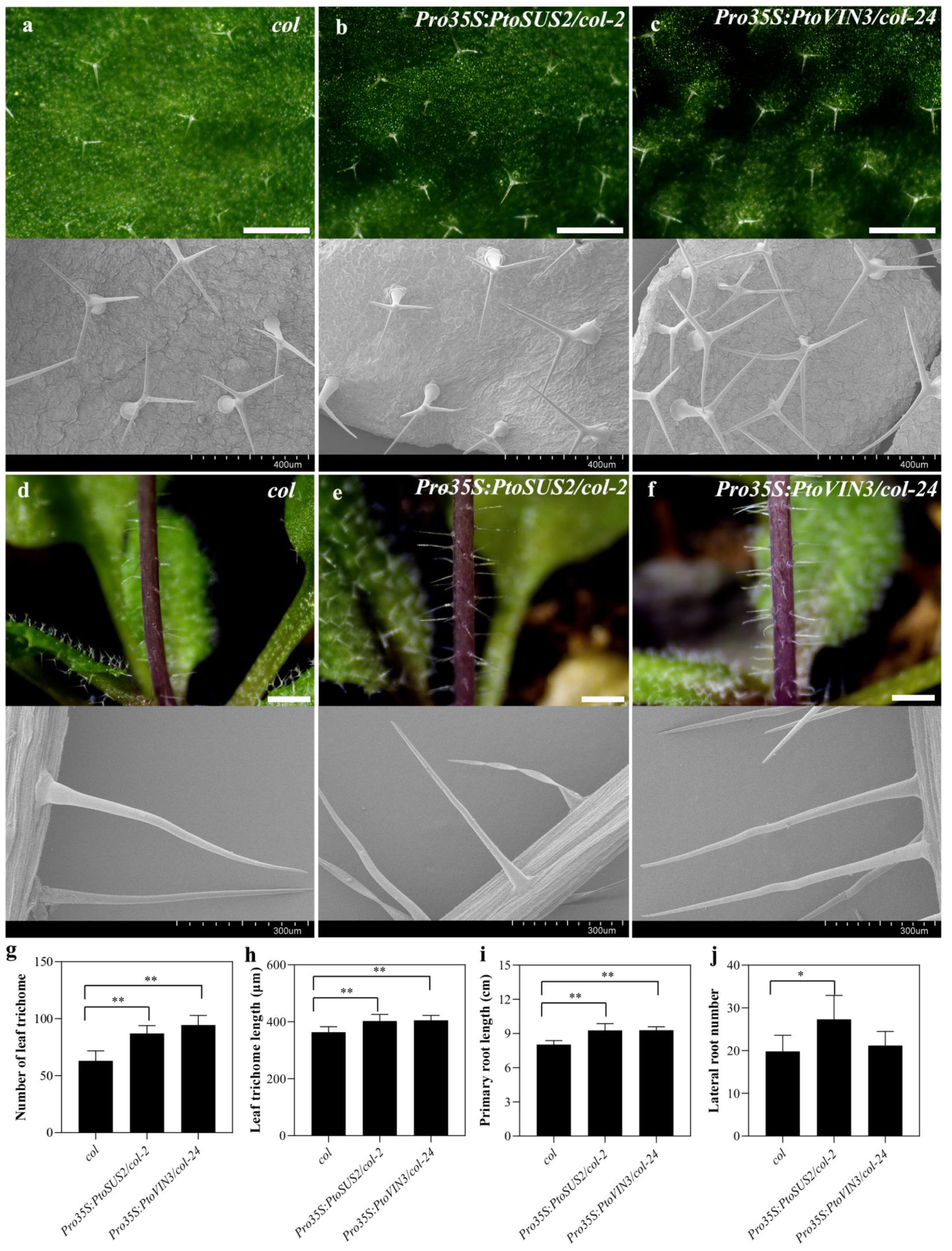
Leaf trichome morphology and phenotypic characteristics in col and its transgenic lines. **a-f** Upper panel, trichome morphology under a stereomicroscope; scale bars = 1 mm. Lower panel, trichome morphology under a scanning electron microscope. **g-l** Phenotypic characteristics of wild-type and transgenic plants; bars represent ± SD, ** significant difference at *P* < 0.01, and * significant difference at *P* < 0.05.

Overexpression of *PtoSUS2* and *PtoVIN3* also led to greater trichome density in mutants *tl1* and *tl2*. Three transgenic lines of each transgenic type were selected for detailed comparison. In *tl1*, eleven and thirteen independent transgenic plants were obtained for *PtoSUS2* and *PtoVIN3*, respectively. Compared with the glabrous phenotype (lacking trichomes) in *tl1*, overexpression of *PtoSUS2* or *PtoVIN3* resulted in normal trichomes on the adaxial surface of rosette leaves; trichomes were observed in the stems of some transgenic plants (Fig. 8a-c; Fig. S8). In *tl2*, twelve and eleven independent transgenic plants were obtained for *PtoSUS2* and *PtoVIN3*, respectively. In *tl2*, trichomes on the adaxial surface of rosette leaves were malformed; few trichomes could be distinguished on the leaf edges. Overexpression of *PtoSUS2* or *PtoVIN3* led to greater malformed trichome density in rosette leaves, especially along the leaf margins. However, the glabrous phenotype of stems was unchanged in *tl2* transgenic plants (Fig. 8d-f; Fig. S9).

**Figure 8.**
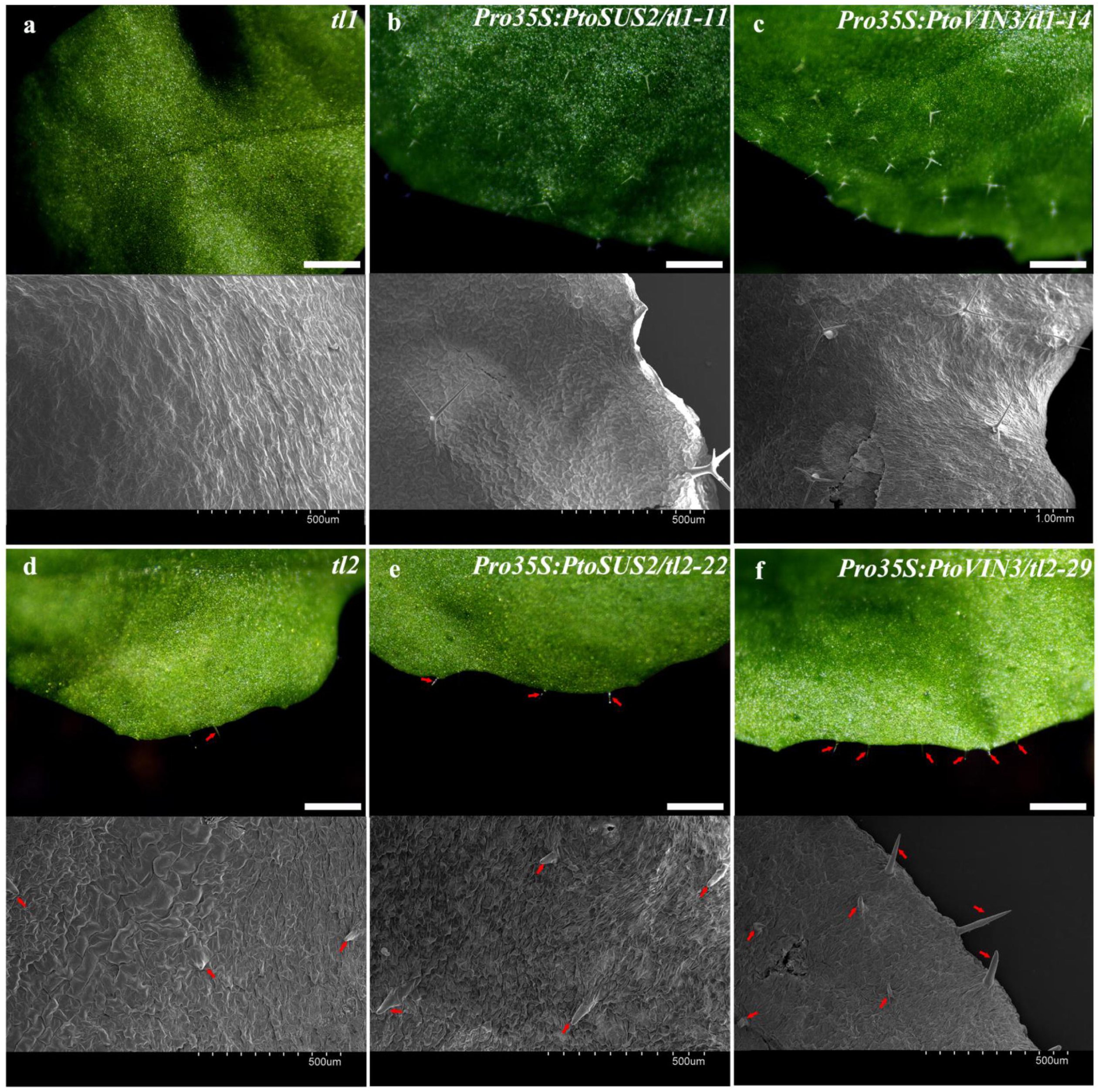
Morphological characteristics of leaf trichome in *tl1/2* and their transgenic lines. Upper panel, trichome morphology under a stereomicroscope; scale bars = 1 mm. Lower panel, trichome morphology under a scanning electron microscope.

### Changes in expression of trichome-related genes in *A. thaliana* **overexpressing *PtoSUS2* and *PtoVIN3***

RT-qPCR was performed to evaluate the expression levels of the exogenous *PtoSUS2* and *PtoVIN3* genes. The two exogenous genes were strongly expressed in the transgenic lines but were undetectable in wild-type *A. thaliana* (Fig. 9). Endogenous genes associated with trichome development were also influenced by *PtoSUS2* and *PtoVIN3*. Overexpression of *PtoSUS2* or *PtoVIN3* in the wild-type increased the expression levels of *AtGL1*, *AtGL2*, *AtGL3*, *AtTTG1*, and *AtMYB23*. The expression of *AtGL1* had a trend similar to the trends of exogenous *PtoSUS2* and *PtoVIN3*; it played a major role in the trichome phenotype of transgenic lines. Similar results were obtained in mutants *tl1* and *tl2*. The phenotype of transgenic lines was closely related to gene expression. For individuals with the most pronounced phenotype in *tl1*, the expression levels of *AtGL2*, *AtGL3*, and *AtMYB23* were increased in *Pro35S:PtoSUS2/tl1-11* and *Pro35S:PtoVIN3/tl1-14*. In *tl2*, elevated expression levels of *PtoSUS2* and *PtoVIN3* led to increased expression of *ATGL1*, *AtGL3*, *AtMYB23*, and *AtTTG1*, especially the results in *Pro35S:PtoSUS2/tl2-22* and *Pro35S:PtoVIN3/tl2-29*. In different transgenic lines, the expression patterns of *AtGL1*, *AtGL3*, and *AtMYB23* showed trends similar to *PtoSUS2* and *PtoVIN3;* they were significantly increased, compared with *tl2*.

**Figure 9.**
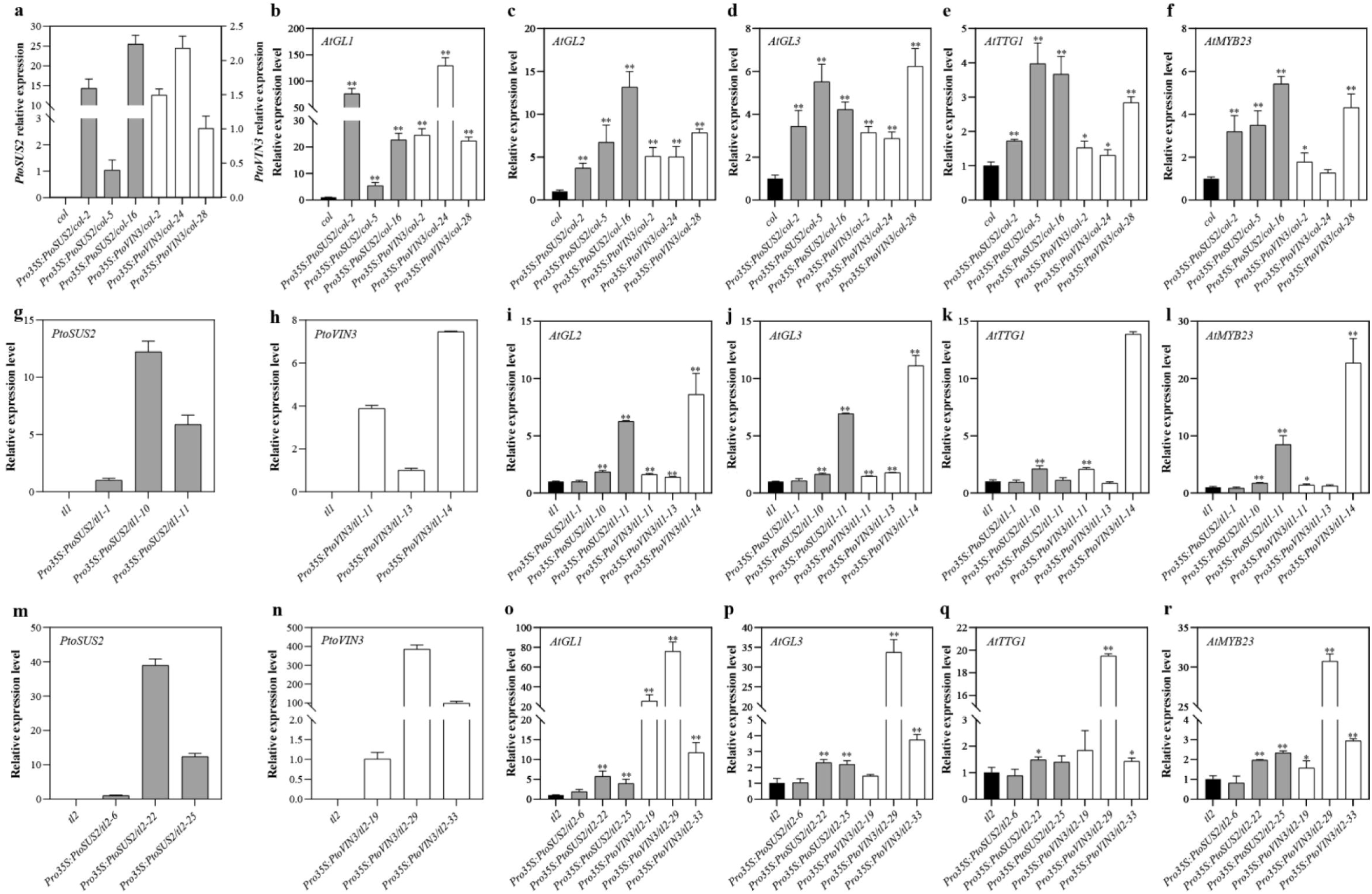
Gene expression patterns in col, tl1/2, and their transgenic lines, as determined RT-qPCR. Bars represent ± SD, ** significant difference at *P* < 0.01, and * significant difference at *P* < 0.05.

## Discussion

Poplar catkin fibers and cotton fibers all belong to plant trichomes, however, there are significant differences in original site. Poplar catkin fibers were not seed hairs and produced at the funicle and placenta (Fig. 2), while cotton fibers were seed fibers and formed from the epidermal cells of fertilized ovule (Ruan *et al*., 2003). In *Arabidopsis*, leaf and stem trichome originate from the epidermal cells of leaves or stems. Root hairs, another type of specific epidermal cells, originate from the epidermal cells of root. Although they all are unicellular structure, there are also differences in morphology. *Arabidopsis* trichome usually have three or four branches, while no branch is found in root hairs, as well as poplar catkin fiber and cotton fiber (Serna and Martin, 2006, Guimil and Dunand, 2007). In addition, root hairs are represented as straight tubular extensions, and performed as position-dependent and unidirectional polarized growth (Roy and Bucksch, 2021, Guimil and Dunand, 2007, Schellmann *et al*., 2002). Poplar catkin fibers consists of hydrophobic microtubes with an external diameter of 8–11 μm and an average length of 0.95–1.59 cm, while the cotton fibers consist of twisted ribbon with an external diameter of 16–20 μm and an average length of 0.95–6.35 cm (Likon *et al*., 2013, Chen and Cluver, 2010). Numerous studies have been done for trichomes, root hairs and cotton fibers (Bui *et al*., 2019, Naoumkina, 2018, Grebe, 2012, Ishida *et al*., 2008), however, less attention has been paid to the poplar catkin fibers. Thus, in this study, the development of poplar catkin fibers was detailed analyzed, and as we performed, sucrose degradation was the primary metabolic factor in catkin fibers.

### Invertase and sucrose synthase have special roles in poplar catkin fibers

In cotton fibers, invertase and sucrose synthase have been shown to play vital roles. Vacuolar invertase, rather than cell wall invertase and neutral/alkaline invertase, had high activity at initiation stage (5 DPA) and then decreased in cotton (Wang *et al*., 2010), but in this study, high and steady activities were both found for vacuolar invertase and neutral/alkaline invertase during poplar catkin fibers development (Fig. 3). In addition, different gene expression patterns of *SUS* were also exhibited in cotton and poplar. Consequently, we speculate that there are special roles for invertase and sucrose synthase during poplar catkin fibers development.

Invertase has a function during the early development of poplar catkin fibers. According to the pH and subcellular localization, invertases can be divided into cell wall invertase, vacuolar invertase and neutral/alkaline invertase, all of which have different functions in plants (Ruan, 2014). Although cell wall invertase and vacuolar invertase both belong to the acid invertase, they can be divided into two subgroups based on evolutionary relationships (Fig. S1a). In cotton fibers, *CWIN* appears to have no regulatory effect in cotton fibers, based on its low and steady expression level (Wang *et al*., 2010, Ruan and Chourey, 1998). Similarly, low cell wall invertase activity was observed in poplar catkin fibers (Fig. 3c). In contrast with cell wall invertase, vacuolar invertase has been proven to participate in both initiation and elongation process in cotton fibers. The hexose produced by *GhVIN1* can be sensed by hexokinase, and then influences cotton fiber initiation though MYB TFs and auxin signaling (Wang *et al*., 2014). Meanwhile, hexose production can promote fiber elongation by increasing osmotic potential (Wang *et al*., 2010). Two other members of *VIN* were shown to be associated with cotton fiber development (Taliercio *et al*., 2010). Among them, *GhVIN2*, probably an allele of *GhVIN1*, was expressed primarily in elongating fibers, and decreased during fiber maturation. Compared with results in cotton fibers, vacuolar invertase activity had a similar pattern during poplar catkin fibers development, indicating that it probably plays the similar role in poplar catkin fibers. Thereinto, *PtoVIN3* has a closest relationship with *GhVIN1* and *GhVIN2*. Surprisingly, neutral/alkaline invertase appeared to have a function during the early development of poplar catkin fibers, differing from cotton fibers. In contrast with their low expression levels in cotton fiber (Wang *et al*., 2010), the neutral/alkaline invertase high activity in poplar catkin fibers (Fig. 3c) led us to infer that neutral/alkaline invertase function differs between poplar and cotton.

Sucrose synthase has a close correlation with the elongation of poplar catkin fibers. In this study, sucrose synthase showed greater activity than invertase (Fig. 3e, f), indicating a potential role in poplar catkin fibers development. In cotton, sucrose synthase has been proposed to play direct role in fiber yield and quality (Ahmed *et al*., 2018). *GhSUS3* was firstly found to have high expression during cotton fiber development, and was thus thought to have a role in that process (Ruan *et al*., 1997). Subsequently, fiberless mutant and sucrose synthase-suppression transgenic plant confirmed the function of *GhSUS3* (Ruan *et al*., 2003, Ruan and Chourey, 1998). In the fiberless mutant and transgenic plant, the activity of sucrose synthase had a close relationship with fiber quality. Furthermore, Elizabeth et al. (Elizabeth *et al*., 2011) found four isoforms of sucrose synthase that are expressed during cotton fiber development. Three of these four members were similar to *GhSUS3*, with high expression in fibers at the extension stage (at 8 DPA). Another isoform, *GhSUSC*, was highly expressed in fibers during secondary cell wall synthesis (at 20 DPA) (Elizabeth *et al*., 2011). Different results were obtained in *Gossypium arboretum* (Chen *et al*., 2012), with four homologous genes having high expression during fiber development (at 15 DPA). However, in poplar, only *PtoSUS2* had high expression at extension stage (stage 4), while the six members in other two clades had consistently low expression (Fig. 4h-j), similar to the pattern in xylem (Zhang *et al*., 2011). High sucrose synthase activity was accompanied by a high hexose to sucrose ratio, which was also observed in cotton fibers (Ruan *et al*., 1997). High turgor pressure, caused by high hexose content, is required for rapid cell extension (Ruan and Chourey, 1998). Therefore, sucrose synthase probably promotes catkin fibers elongation by breaking down sucrose to increase turgor pressure. Although vacuolar invertase activity was also high during catkin fibers extension, fitting analysis suggested that sucrose synthase plays the dominant role (Fig. 3).

### Specific regulatory mechanisms during poplar catkin fibers development

To further elucidate the role of sucrose in poplar catkin fibers, we investigated TFs and other genes that may be involved in the development process, and co-expression network and YIH assay were performed (Fig. 5h, i, Fig. 6, Table S1). In *Arabidopsis*, a module of regulatory genes, *GL1*/*GL3*/*TTG1*-*GL2*, have been proven to play a key regulatory role in the development of trichomes in many researches (review in (Bui *et al*., 2019)). *WEREWOLF*, the homologous of *GL1*, worked together with *GL3*, *TTG1* and *GL2* to regulate hairless cell differentiation in root (Ishida *et al*., 2008). Similar regulatory patterns were also found in cotton fibers. *GhMYB109* (Suo *et al*., 2003), *GhDEL65* (Shangguan *et al*., 2016) and *GhTTG1/3* (Humphries *et al*., 2005) are homologues of *GL1* (MYB), *GL3/EGL3* (bHLH) and *TTG1* (WD40), respectively, in cotton and are thought to exert effects on fiber initiation. *Pto(GL1)-L1*, *Pto(GL3)-L1* and *2* and *Pto(TTG1)-L1*, which are the representatives in poplar, all showed increased expression at the initiation stage of catkin fibers development. *GhMYB25* and its homologous, another type of MYB TFs, was found to play regulatory roles in fiber initiation and extension (Wu *et al*., 2018, Walford *et al*., 2011, Machado *et al*., 2009), however their homologues had delayed expression in poplar catkin fibers, aside from *Pto(GhMYB25)-L1/2* (Fig. 5h). In addition, previous studies about cotton have shown that sucrose transpotation regulated by *GhMYB212* has an effect on fiber elongation (Sun *et al*., 2019), and *GhVIN1*-derivied hexose signaling has potential regulated role on the key MYB TFs, including *GhMYB25*, *GhMYB25-like* and *GhMYB109* (Wang *et al*., 2014). Although the roles of sucrose metabolism in trichomes development has been confirmed by many researches in cotton (Naoumkina, 2018), the relationship between sucrose metabolism and key TFs, such as *GL1*, *GL2*, *GL3* and so on, is still unclear. In this study, co-expression network showed only *Pto(GL3)-L1/2* has a close relationship with sucrose metabolism related genes (cluster I, Fig. 6a). As the key roles of sucrose synthase and vacuolar invertase in poplar catkin fibers development, combining the results of RT-qPCR and correlation analysis, two key genes, *PtoSUS2* and *PtoVIN3* were selected for in-depth analysis. Interestingly, analysis of promoter regions and Y1H showed that MYB and bHLH TFs maybe the potential regulators of *PtoSUS2*/*PtoVIN3* (Fig. 6c-d). Notably, *Pto(GhMYB25)-L2/4,* the homologue of *Populus MYB186* and *MYB38*, had a strong interaction with *Pro-PtoSUS2* and slightly interaction with *Pro-PtoVIN3*. Recent study showed that positive leaf and stem trichome regulator *MYB186* and its paralog *MYB38* is also essential for seed trichome development in hybrid poplar (Patrick Bewg *et al*., 2022, Plett *et al*., 2010, Ortega *et al*., 2022), which indicated that *PtoSUS2* and *PtoVIN3* maybe the potential targeting genes of trichome-regulating MYB and were regulated by MYBs to affect the development of poplar catkin fibers.

In addition, *CesA* genes also play different roles between poplar catkin fibers and cotton fiber. The *CesA* gene family has been found to play roles during cotton fiber development (Haigler *et al*., 2009). To elaborate, the global genome analysis of the *CesA* gene family in cotton was performed (Li *et al*., 2013a). Members of the S1, S2 and S3 clades showed preferential expression during secondary cell wall synthesis, while other members appeared to play a role in cotton fiber initiation and extension. Similar to cotton (Li *et al*., 2013a, Zhang *et al*., 2012), the gradually increased cellulose content (Fig. S2) and negatively correlation between cellulose and sucrose (Table 1) also suggest that cellulose plays an important role in poplar catkin fibers. However, cellulose synthase seems to have a different role in poplar catkin fibers. Compared with the secondary cell wall synthesis-related members, members involving in primary cell wall synthesis played a more critical role as their high expression (Fig. 5a-g). And compared with the homologous in cotton (Li *et al*., 2013a), less changes of cellulose synthase genes expression were observed in poplar catkin fibers, and we speculate that this difference underlies the different qualities of catkin fibers in the two plants. 44.5% cellulose content in poplar catkin fibers was determined by α-cellulose, while more than 80% of cellulose in cotton fibers was β-cellulose (Zhang *et al*., 2018, Murugesh Babu *et al*., 2013). Correlation analysis showed that cellulose content has a significant reverse correlation with sucrose content. Similar results were also found in cotton. Cellulose content increased rapidly and sucrose content decreased rapidly from 17 DPA to 45 DPA in cotton fibers (Ma *et al*., 2014). Further research in cotton have showed that the sucrose content is one of the main reasons for the difference in cellulose content in different varieties (Zhang *et al*., 2012). These results also confirmed that the sucrose metabolism process is important to the fiber development process, such as poplar catkin fibers.

### *PtoSUS2* and *PtoVIN3* increased trichome density in *A. thaliana*

In poplar, *PtoSUS2* is widely expressed in roots, vegetative buds, and floral catkins; its homologue *PtrSUS2* is highly expressed in xylem tissues and moderately expressed in leaves and roots (An *et al*., 2014, Zhang *et al*., 2011). In this study, *PtoSUS2* had higher expression than other members of the *SUS* gene family during poplar catkin fibers development; it was significantly correlated with sucrose synthase activity (Fig. 4, Table 1). Therefore, *PtoSUS2* may play a major role in poplar catkin fibers development; this hypothesis was supported by the results of *PtoSUS2* overexpression in *A. thaliana*. Overexpression wild-type plants had increased trichome density on rosette leaves and stems, and transgenic plants of the glabrous mutants produced more trichomes on rosette leaves (Fig. 7, Fig. 8). *GhSUS3*, which has a close relationship with *PtoSUS2* in cotton, is implicated in cotton fiber cell initiation and elongation (Ruan *et al*., 2003). Overexpression of *SUS* increases cotton fiber length, while decreased expression of *GhSUS3* induces the fiberless mutation (Ahmed *et al*., 2018, Ruan and Chourey, 1998). Similar results were also found for *PtoSUS2*. Ectopic expression of *PtoSUS2* not only increased the leaf trichome density, but also increased the trichome length, which indicated that *PtoSUS2* plays dual roles in the trichome initiation and elongation. Another candidate gene, *PtoVIN3*, was constitutively expressed in roots and leaves; it is also regulated by abiotic and biotic stresses (Chen *et al*., 2015). In this study, overexpression of *PtoVIN3* also increased trichome density and length on rosette leaves, indicating that it functions with *PtoSUS2* to control catkin fibers development in poplar. Similar results have also been proved in cotton. High vacuolar invertase activity is important for cotton fiber development; the homologue of *PtoVIN3*, *GhVIN1*, is a key factor in that process (Wang *et al*., 2010). Notably, the mutants of *PtoSUS2* and *PtoVIN3* homologous in *Arabidopsis* did not obtained fiberless phenotype, and the increased density of trichomes is not due to enhanced vegetative growth in *A. thaliana* transgenic lines (Fig. S6). However, the influence on trichome density of both rosette leaves and stems, and primary root length showed that the specific differentiation of epidermal cells is influenced at the whole plant level.

Trichome development in *A. thaliana* is controlled by the *AtGL1*-*AtGL3*-*AtTTG1* complex; *AtGL2* acts downstream of this complex to control normal trichome development (Pesch *et al*., 2015). *AtMYB23* controls trichome branching and initiation at leaf edges; it is functionally redundant with *AtGL1* (Kirik *et al*., 2005). In wild-type plants overexpressing *PtoSUS2* and *PtoVIN3*, the expression levels of all above-mentioned genes were increased. This demonstrates the effect of *PtoSUS2* and *PtoVIN3* on trichome development in *A. thaliana* at the molecular level. In different transgenic lines, the expression patterns of *AtGL1* are closely like the expression of *PtoSUS2* or *PtoVIN3*, which was consistent with our Y1H assay results. *PtoSUS2* and *PtoVIN3* have a close relationship with *Pto(GhMYB25)-L2/4*, the homologous of *AtGL1*. Fiberless mutants used in our study, *tl1* and *tl2*, were the mutants of *GL1* and *GL2*, respectively. Over-expression of *PtoSUS2* and *PtoVIN3* partially restored the trichome phenotype in the two muntants, which also indirectly confirmed the relationship between sucrose metabolism and the key TFs during trichome development. Additionally, *GhVIN1* regulates fiber initiation from the ovule epidermis, probably via sugar signaling with *cis*-acting elements of the sugar-response element in *GhMYB25* and *GhMYB25like* (Wang *et al*., 2014), which is similar to poplar catkin fibers development. The promoter regions of *Pto(GL1)-L1/2/3* and *Pto(GhMYB25)-L1/2/3/4*, the homologues of *AtGL1* (in *Arabidopsis*)and *GhMYB25* (in cotton), have also been analyzed in poplar. SURE motifs (i.e., sugar-responsive elements) (Wang *et al*., 2014), were found in the promoter region of *Pto(GhMYB25)-L1/2/4* (Fig. S10). Additionally, the SP8B-like motif (TACT(X)TT), which is similar to another sugar-responsive element (Wang *et al*., 2014), was found in the promoter regions of *Pto(GL1)-L2* and *Pto(GhMYB25)-L3/4*. Therefore, we speculate that *PtoSUS2* and *PtoVIN3* probably regulate catkin fibers development via sugar signaling.

In this study, sucrose metabolism was analyzed in detail in the context of poplar catkin fibers development, and sucrose degradation was found to play the dominant role in the process. As shown in Fig. 10, sucrose is consumed rapidly as poplar catkin fibers mature, especially at the extension stage. Vacuolar invertase and sucrose synthase, probably derived by *PtoVIN3* and *PtoSUS2*, which were the potential targeting genes of trichome-regulating MYBs, were thought to perform the function. Furthermore, the PCW synthesis-related *PtoCesAs* may play a major role in cellulose synthesis. As an important metabolic process in poplar catkin fibers, analyzing the regulatory mechanism of sucrose metabolism is not only helpful for us to understand the regulatory mechanism in woody plants, and for identifying key candidate genes with high breeding value in *P. tomentosa*, but also provides a foundation for solving environmental problems caused by poplar catkins though genetic engineering.

**Figure 10.**
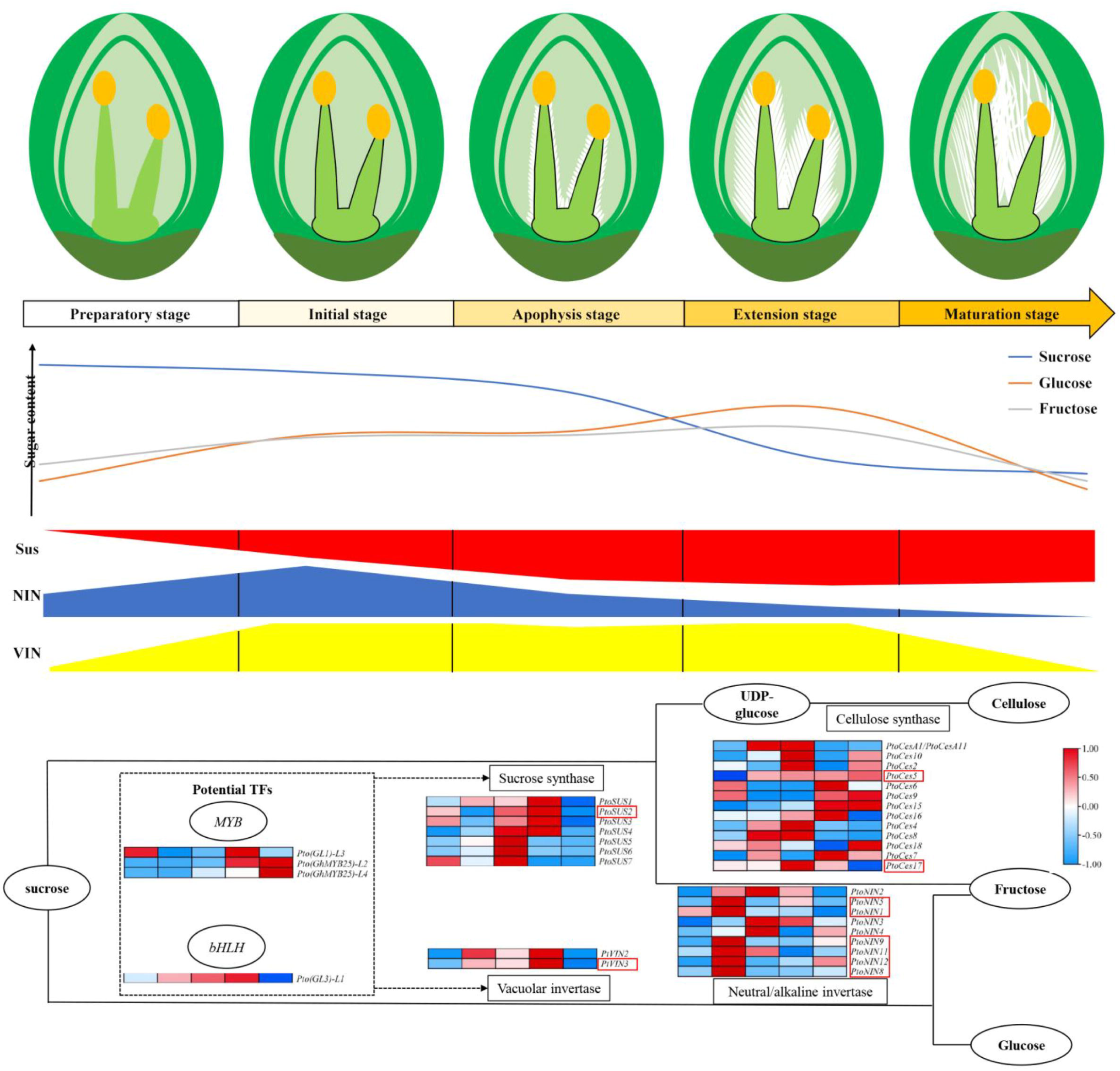
Potential regulatory model of sucrose metabolism in poplar catkin fibers. First panel represented morphological diagram of poplar catkin fibers. Second panel represented the change patterns of sucrose, glucose and fructose. Third panel represented the enzyme activity of sucrose synthase, invertase and vacuolar invertase. Last panel represented expression patterns of *SUS*, *NIN*, *VIN*, *CesA*, potential *MYB* and *bHLH* TFs, and the genes in the red box are key candidate genes as their high expression level and close relationship with sugar and cellulose contents.

## Supplementary data

**Figure S1 Phylogenetic tree of the sucrose metabolism-related gene families.**

**Figure S2 The cellulose content in poplar catkin fibers.**

**Figure S3 Analysis of cis-acting elements for sucrose metabolism-related genes in *P. tomentosa*.**

**Figure S4 Tissue-specific expression analysis of *PtoSUS2* and *PtoVIN3*.**

**Figure S5 Morphological characteristics of trichome on rosette leaves and stems in *A. thaliana* wild-type and transgenic lines.**

**Figure S6 Characteristics of root, rosette leaves, and whole plant in *A. thaliana* wild-type and transgenic lines.**

**Figure S7 Changed in trichome density on leaves during leaf development in *A. thaliana* wild-type and transgenic lines.**

**Figure S8 Morphological characteristics of trichome on leaves and stems in transgenic lines of *tl1* and *tl1*.**

**Figure S9 Morphological characteristics of trichome on leaves and stems in transgenic lines of *tl2* and *tl2*.**

**Figure S10 Analysis of sugar signal-related cis-acting elements for *Pto(GL1)-L1/2/3* and *Pto(GhMYB25)-L1/2/3/4*.**

**Table S1 Comparison of potential TFs and other genes in poplar and cotton**

**Table S2 Primers used for RT-qPCR**

**Table S3 Accession numbers of genes used for phylogenetic tree construction**

**Table S4 Comparison of the TFs for YIH in *poplar* and *Arabidopsis***

**Table S5 Primers used for plasmid construction**

**Methods S1: Quantitation of sugar content and enzyme activities Methods S2: Quantitation of cellulose content**

## Acknowledgements

We gratefully acknowledge the families who participated by giving research assistance. We also thank the families who participated by giving research assistance.

## Author contributions

Z.C. and X.Y. designed the work; X.Y., T.Z. and N.Y. performed experiments and analyzed data; Co-expression network analysis was carried out by P.R.; X.Y. and T. Z. drafted the manuscript; and Z.C., G.L., L. J. and X.A. revised the manuscript. X. Y. and T. Z. contributed equally to this work.

## Data availability

The data supporting the findings of this study are available from the corresponding author, Zhong Chen, upon request.

## Abbreviations

CesA: cellulose synthase
Col: Columbia ecotype
CWIN: cell wall invertase
DAG: days after germination
EGL3: ENHANCER OF GLABRA3
GL1: GLABROUS1
GL2: GLABRA2
GL3: GLABROUS3
NIN: neutral/alkaline invertase
SPS: sucrose phosphate synthase
SPP: sucrose phosphate phosphatase
SUS: sucrose synthase
TF: transcription factor
tl1: trichome less 1
tl2: trichome less 2
TTG1: TRANSPARENT TESTA GLABRA1
VIN: vacuolar invertase

## References

Ahmed M, Shahid AA, Akhtar S, Latif A, Ud Din S, Fanglu M, et al. 2018. Sucrose synthase genes: a way forward for cotton fiber improvement. Biologia, 73, 703–713.

An X, Chen Z, Wang J, Ye M & Ji L 2014. Identification and characterization of the Populus sucrose synthase gene family. Gene Amsterdam.

An X, Gao K, Chen Z, Li J, Yang X, Yang X, et al. 2022. High quality haplotype-resolved genome assemblies of Populus tomentosa Carr., a stabilized interspecific hybrid species widespread in Asia. Mol Ecol Resour, 22, 786–802.

An XW, Dongmei; Wang, Zeliang; Wang, Jingcheng; Cao, Guanlin; Bo, Wenhao; Zhang, Zhiyi 2010. Expression profile of PtLFY in floral bud development associated with floral bud morphological differentiation in Populus tomentosa. Sci Silvae Sinicae, 46, 32–38.

Bui APN, Ong B-N, Huu T-aN & Tran H-D 2019. Similarities between Initiation Mechanism between Cotton Fiber and Arabidopsis Trichome: Prospects in Improving Cotton Fiber Yield. annual research & review in biology, 1–9.

Chen A, He S, Li F, Li Z, Ding M, Liu Q, et al. 2012. Analyses of the sucrose synthase gene family in cotton: structure, phylogeny and expression patterns. BMC plant biology, 12, 85.

Chen HL & Cluver R 2010. Assessment of Poplar Seed Hair Fibers as a Potential Bulk Textile Thermal Insulation Material. Clothing & Textiles Research Journal, 28, p.255–262.

Chen Z, Gao K, Su X, Rao P & An X 2015. Genome-Wide Identification of the Invertase Gene Family in Populus. plos one, 10.

Chen Z, Rao P, Yang X, Su X, Zhao T, Gao K, et al. 2018. A Global View of Transcriptome Dynamics During Male Floral Bud Development in Populus tomentosa. Scientific Reports, 8, 722.

Elizabeth B, Michel VT, G WR, Danny L, M CP, Steven E, et al. 2011. A novel isoform of sucrose synthase is targeted to the cell wall during secondary cell wall synthesis in cotton fiber. Plant Physiology, 157, 40–54.

Fujii S, Hayashi T & Mizuno K 2010. Sucrose synthase is an integral component of the cellulose synthesis machinery. Plant and cell physiology, 51, 294–301.

Grebe M 2012. The patterning of epidermal hairs in Arabidopsis : updated. current opinion in plant biology, 15, 31–37.

Guimil S & Dunand C 2007. Cell growth and differentiation in Arabidopsis epidermal cells. Journal of Experimental Botany, 58, 3829–3840.

Haigler CH, Singh B, Wang G & Zhang D 2009. Genomics of cotton fiber secondary wall deposition and cellulose biogenesis. In: Paterson, A. H. (ed.) Genetics and genomics of cotton. New York, NY: Springer.

Haigler CH, Singh B, Zhang D, Hwang S, Wu C, Cai WX, et al. 2007. Transgenic cotton over-producing spinach sucrose phosphate synthase showed enhanced leaf sucrose synthesis and improved fiber quality under controlled environmental conditions. Plant Molecular Biology, 63, 815–832.

Humphries JA, Walker AR, Timmis JN & Orford SJ 2005. Two WD-repeat genes from cotton are functional homologues of the Arabidopsis thalianaTRANSPARENT TESTA GLABRA1 (TTG1) gene. Plant molecular biology, 57, 67–81.

Ishida T, Kurata T, Okada K & Wada T 2008. A Genetic Regulatory Network in the Development of Trichomes and Root Hairs. Annual Review of Plant Biology, 59, 365–386.

Kirik V, Lee MM, Wester K, Herrmann U, Zheng Z, Oppenheimer D, et al. 2005. Functional diversification of MYB23 and GL1 genes in trichome morphogenesis and initiation. Development, 132, 1477–1485.

Klocko AL, Brunner AM, Huang J, Meilan R, Lu H, Ma C, et al. 2016. Containment of transgenic trees by suppression of LEAFY. Nat Biotechnol, 34, 918–22.

Li A, Xia T, Xu W, Chen T, Li X, Fan J, et al. 2013a. An integrative analysis of four CESA isoforms specific for fiber cellulose production between Gossypium hirsutum and Gossypium barbadense. Planta, 237, 1585–1597.

Li B, Li D-D, Zhang J, Xia H, Wang X-L, Li Y, et al. 2013b. Cotton AnnGh3 Encoding an Annexin Protein is Preferentially Expressed in Fibers and Promotes Initiation and Elongation of Leaf Trichomes in Transgenic Arabidopsis. Journal of Integrative Plant Biology, 55, 902–916.

Likon M, Remškar M, Ducman V & Švegl F 2013. Populus seed fibers as a natural source for production of oil super absorbents. Journal of Environmental Management, 114, 158–167.

Lu H, Klocko AL, Brunner AM, Ma C, Magnuson AC, Howe GT, et al. 2019. RNA interference suppression of AGAMOUS and SEEDSTICK alters floral organ identity and impairs floral organ determinacy, ovule differentiation, and seed-hair development in Populus. New Phytol, 222, 923–937.

Ma Y, Wang Y, Liu J, Lv F, Chen J & Zhou Z 2014. The effects of fruiting positions on cellulose synthesis and sucrose metabolism during cotton (Gossypium hirsutum L.) fiber development. PLoS One, 9, e89476.

Machado A, Wu Y, Yang Y, Llewellyn DJ & Dennis ES 2009. The MYB transcription factor GhMYB25 regulates early fibre and trichome development. The Plant Journal, 59, 52–62.

Martin LK & Haigler CH 2004. Cool temperature hinders flux from glucose to sucrose during cellulose synthesis in secondary wall stage cotton fibers. Cellulose, 11, 339–349.

Müller NA, Kersten B, Leite Montalvão AP, Mähler N, Bernhardsson C, Bräutigam K, et al. 2020. A single gene underlies the dynamic evolution of poplar sex determination. Nat Plants, 6, 630–637.

Murugesh Babu K, Selvadass M & Somashekar R 2013. Characterization of the conventional and organic cotton fibres. The Journal of The Textile Institute, 104, 1101–1112.

Naoumkina M 2018. Advances in Understanding of Cotton Fiber Cell Differentiation and Elongation. In: Fang, D. D. (ed.) Cotton Fiber: Physics, Chemistry and Biology. Cham: Springer International Publishing.

Ortega MA, Zhou R, Chen MSS, Bewg WP, Simon B & Tsai C-J 2022. In vitro floral development in poplar: insights into seed trichome regulation and trimonoecy. New Phytologist, n/a.

Park JY, Canam T, Kang KY, Ellis DD & Mansfield SD 2008. Over-expression of an arabidopsis family A sucrose phosphate synthase (SPS) gene alters plant growth and fibre development. Transgenic Research, 17, 181–192.

Patrick Bewg W, Harding SA, Engle NL, Vaidya BN, Zhou R, Reeves J, et al. 2022. Multiplex knockout of trichome-regulating MYB duplicates in hybrid poplar using a single gRNA. Plant Physiol.

Pattanaik S, Patra B, Singh SK & Yuan L 2014. An overview of the gene regulatory network controlling trichome development in the model plant, Arabidopsis. frontiers in plant science, 5, 259–259.

Pesch M, Schultheiß I, Klopffleisch K, Uhrig JF, Koegl M, Clemen CS, et al. 2015. TRANSPARENT TESTA GLABRA1 and GLABRA1 Compete for Binding to GLABRA3 in Arabidopsis. Plant Physiol, 168, 584–97.

Plett JM, Wilkins O, Campbell MM, Ralph SG & Regan S 2010. Endogenous overexpression of Populus MYB186 increases trichome density, improves insect pest resistance, and impacts plant growth. Plant J, 64, 419–32.

Roy A & Bucksch A 2021. Root hairs vs. trichomes: Not everyone is straight! Current Opinion in Plant Biology, 64, 102151.

Ruan Y-L 2014. Sucrose metabolism: gateway to diverse carbon use and sugar signaling. Annual review of plant biology, 65, 33–67.

Ruan Y-L & Chourey PS 1998. A fiberless seed mutation in cotton is associated with lack of fiber cell initiation in ovule epidermis and alterations in sucrose synthase expression and carbon partitioning in developing seeds. Plant physiology, 118, 399–406.

Ruan Y-L, Chourey PS, Delmer DP & Perez-Grau L 1997. The differential expression of sucrose synthase in relation to diverse patterns of carbon partitioning in developing cotton seed. Plant Physiology, 115, 375–385.

Ruan Y-L, Llewellyn DJ & Furbank RT 2003. Suppression of sucrose synthase gene expression represses cotton fiber cell initiation, elongation, and seed development. The Plant Cell, 15, 952–964.

Sannigrahi P, Ragauskas AJ & Tuskan GA 2010. Poplar as a feedstock for biofuels: A review of compositional characteristics. biofuels bioproducts and biorefining, 4, 209–226.

Schellmann S, Schnittger A, Kirik V, Wada T, Okada K, Beermann A, et al. 2002. TRIPTYCHON and CAPRICE mediate lateral inhibition during trichome and root hair patterning in Arabidopsis. The EMBO Journal, 21, 5036–5046.

Serna L & Martin C 2006. Trichomes: different regulatory networks lead to convergent structures. Trends in Plant Science, 11, 274–280.

Shangguan XX, Yang CQ, Zhang XF & Wang LJ 2016. Functional characterization of a basic Helix-Loop - Helix (bHLH) transcription factor GhDEL65 from cotton (Gossypium hirsutum). Physiologia plantarum, 158, 200-212.

Sun W, Gao Z, Wang J, Huang Y, Chen Y, Li J, et al. 2019. Cotton fiber elongation requires the transcription factor Gh MYB 212 to regulate sucrose transportation into expanding fibers. New Phytologist, 222, 864–881.

Suo J, Liang X, Pu L, Zhang Y & Xue Y 2003. Identification of GhMYB109 encoding a R2R3 MYB transcription factor that expressed specifically in fiber initials and elongating fibers of cotton (Gossypium hirsutum L.). Biochimica et Biophysica Acta (BBA)-Gene Structure and Expression, 1630, 25–34.

Taliercio E, Scheffler J & Scheffler B 2010. Characterization of two cotton (Gossypium hirsutum L) invertase genes. Molecular biology reports, 37, 3915–3920.

Walford SA, Wu Y, Llewellyn DJ & Dennis ES 2011. GhMYB25-like: a key factor in early cotton fibre development. The Plant Journal, 65, 785–797.

Wang L, Cook A, Patrick JW, Chen XY & Ruan YL 2014. Silencing the vacuolar invertase gene GhVIN 1 blocks cotton fiber initiation from the ovule epidermis, probably by suppressing a cohort of regulatory genes via sugar signaling. The Plant Journal, 78, 686–696.

Wang L, Li X-R, Lian H, Ni D-A, He Y-K, Chen X-Y, et al. 2010. Evidence that high activity of vacuolar invertase is required for cotton fiber and Arabidopsis root elongation through osmotic dependent and independent pathways, respectively. Plant Physiology, 154, 744–756.

Wu H, Tian Y, Wan Q, Fang L, Guan X, Chen J, et al. 2018. Genetics and evolution of MIXTA genes regulating cotton lint fiber development. new phytologist, 217, 883–895.

Xue L, Wu H, Chen Y, Li X, Hou J, Lu J, et al. 2020. Evidences for a role of two Y-specific genes in sex determination in Populus deltoides. Nat Commun, 11, 5893.

Yang W, Wang D, Li Y, Zhang Z, Tong S, Li M, et al. 2021. A General Model to Explain Repeated Turnovers of Sex Determination in the Salicaceae. Mol Biol Evol, 38, 968–980.

Yang Z, Wang T, Wang H, Huang X, Qin Y & Hu G 2013. Patterns of enzyme activities and gene expressions in sucrose metabolism in relation to sugar accumulation and composition in the aril of Litchi chinensis Sonn. Journal of Plant Physiology, 170, 731–740.

Ye M, Zhong C, Su X, Ji L, Jia W & Liao… W 2014. Study of seed hair growth in Populus tomentosa, an important character of female floral bud development. BMC Genomics, 15.

Zhang D, Xu B, Yang X, Zhang Z & Li B 2011. The sucrose synthase gene family in Populus: structure, expression, and evolution. Tree genetics & genomes, 7, 443–456.

Zhang F, Zuo K, Zhang J, Liu X, Zhang L, Sun X, et al. 2010. An L1 box binding protein, GbML1, interacts with GbMYB25 to control cotton fibre development. Journal of experimental botany, 61, 3599–3613.

Zhang M, Song X, Sun X, Wang Z, Li Z, Ji H, et al. 2012. The relationship between cellulose content and the contents of sugars and minerals during fiber development in colored cotton cultivars. Cellulose, 19, 2003–2014.

Zhang X, Henriques R, Lin SS, Niu QW & Chua NH 2006. Agrobacterium-mediated transformation of Arabidopsis thaliana using the floral dip method. Nature Protocols, 1, 641.

Zhang X, Li Z, Yu Y & Wang H 2018. Characterizations of poplar catkin fibers and their potential for enzymatic hydrolysis. JOURNAL OF WOOD SCIENCE, 64, 458–462.

