## Supplementary figures and images for "Sucrose metabolism and candidate genes during catkin fibers development in poplar"

### Figure S1.tif

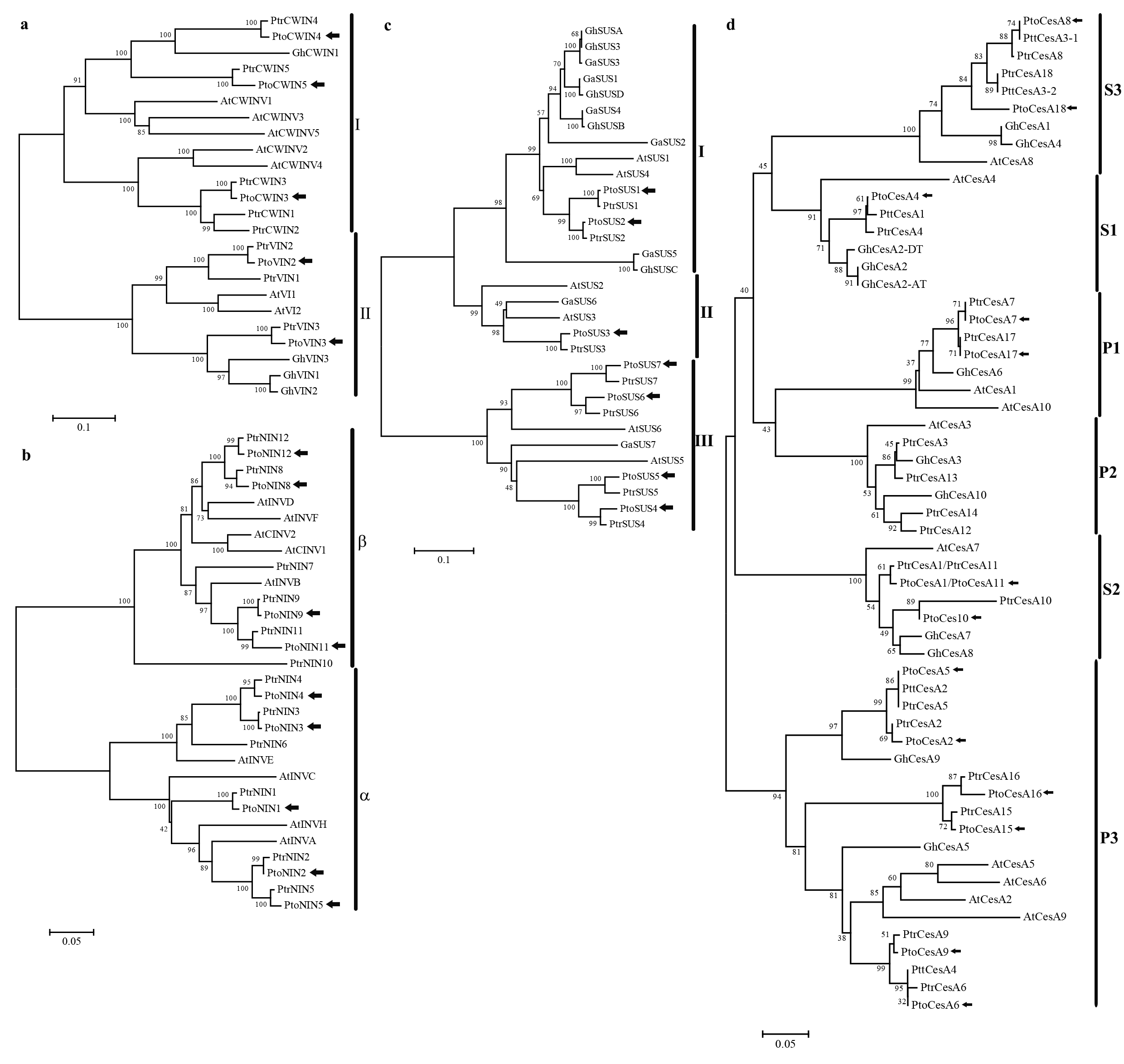

### Figure S2.tif

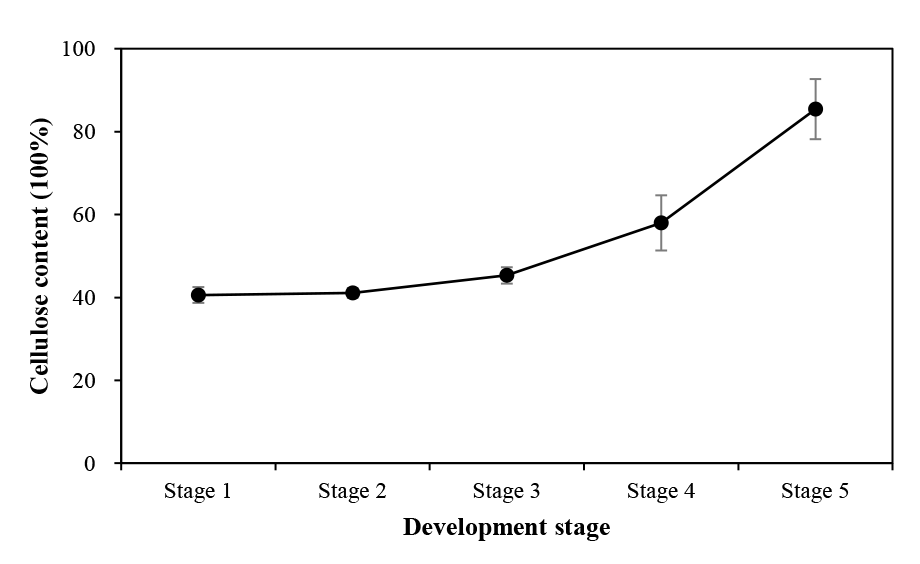

### Figure S3.tif

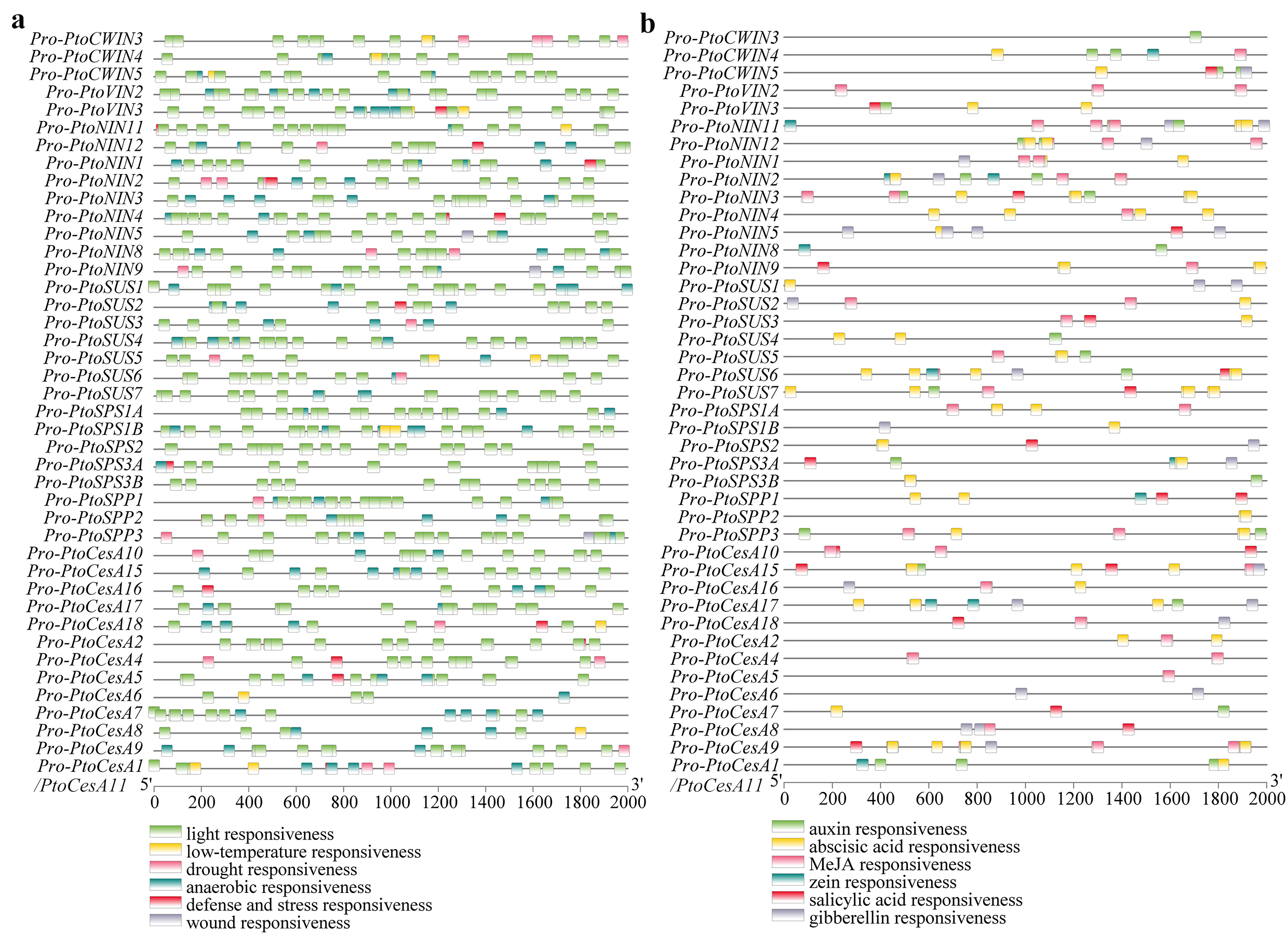

### Figure S4.tif

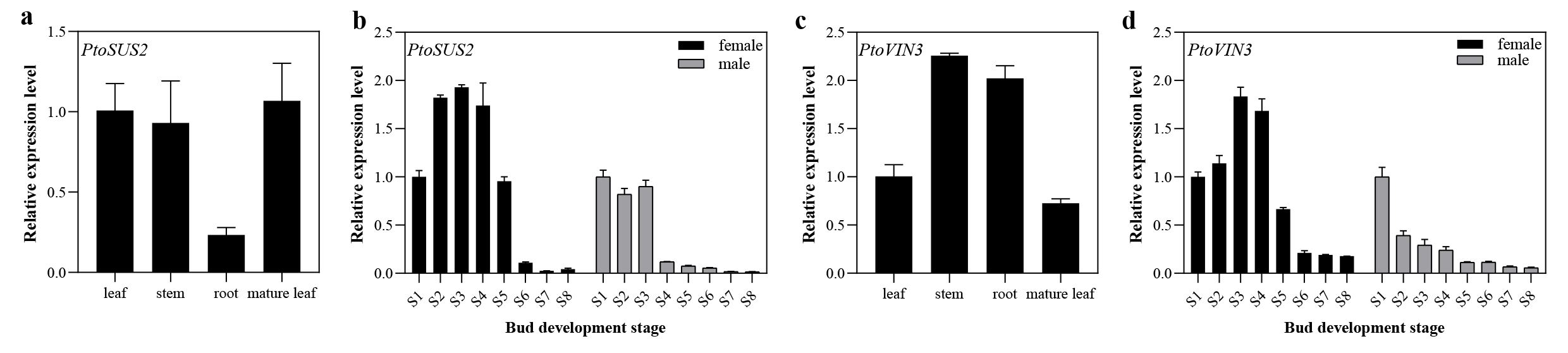

### Figure S4.tif

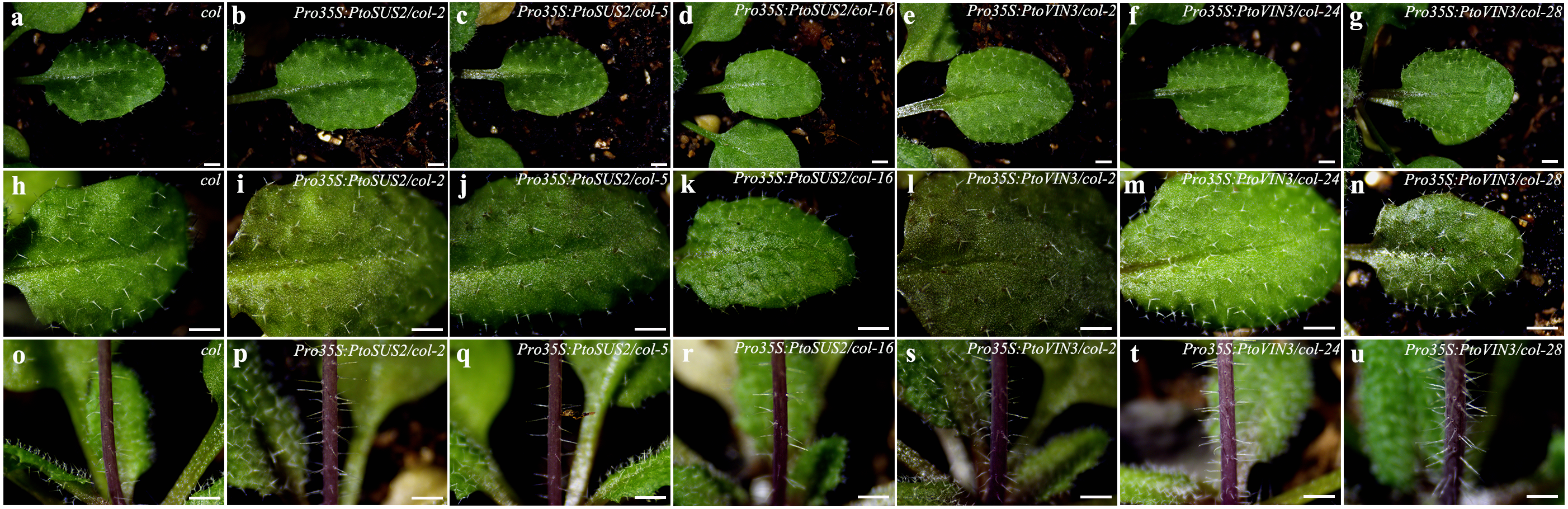

### Figure S6.tif

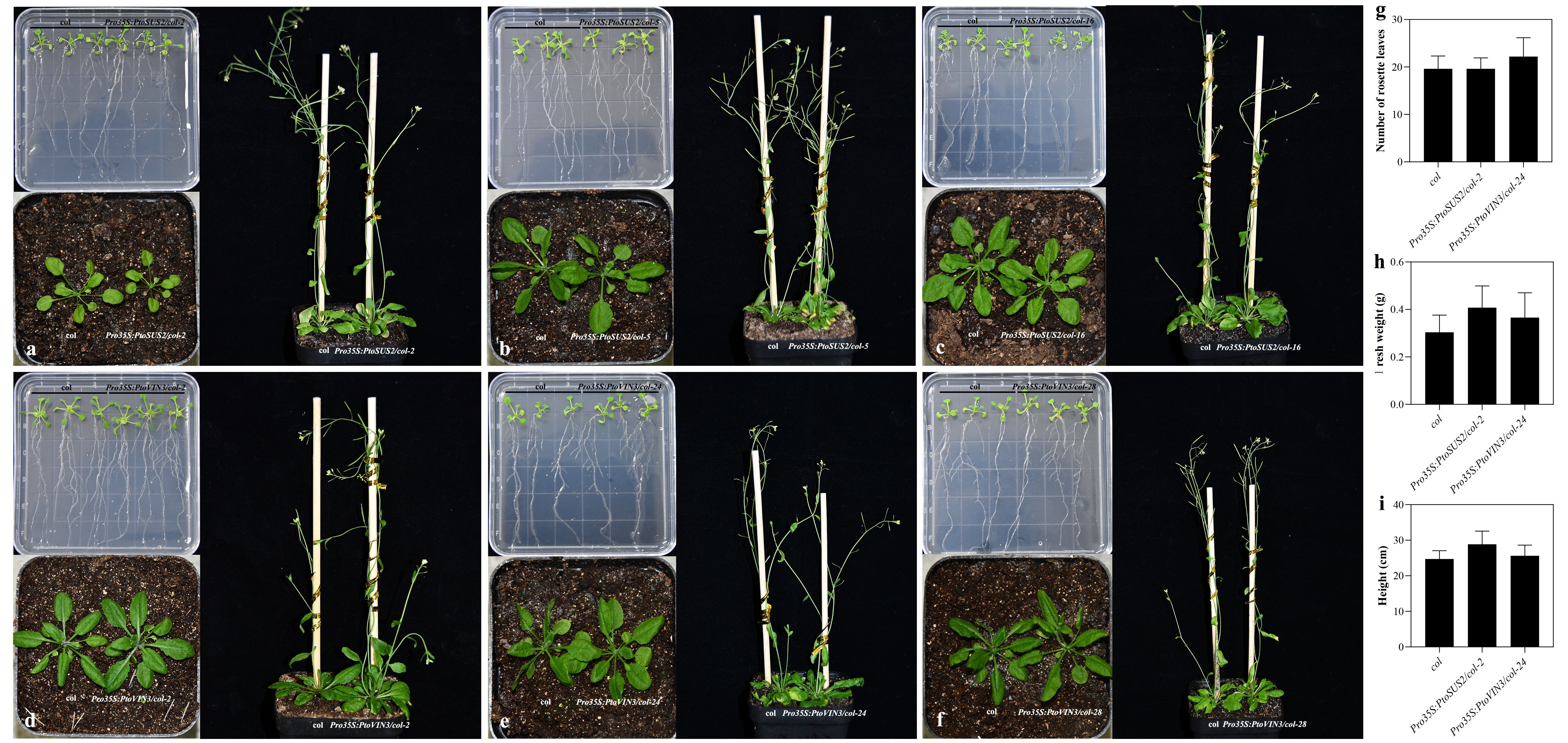

### Figure S7.tif

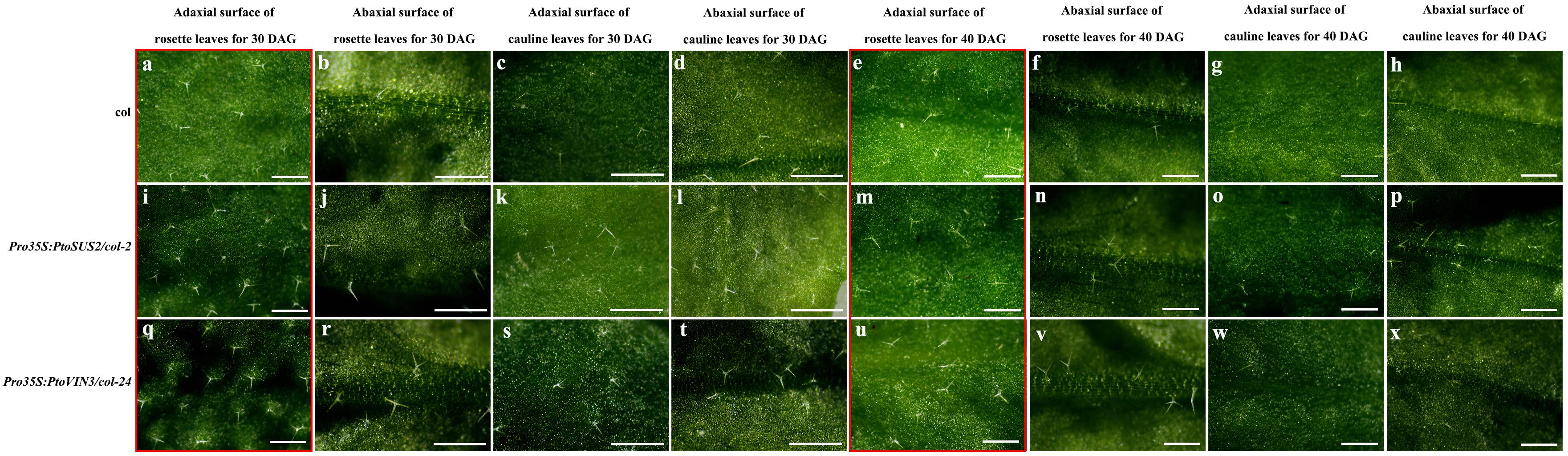

### Figure S8.tif

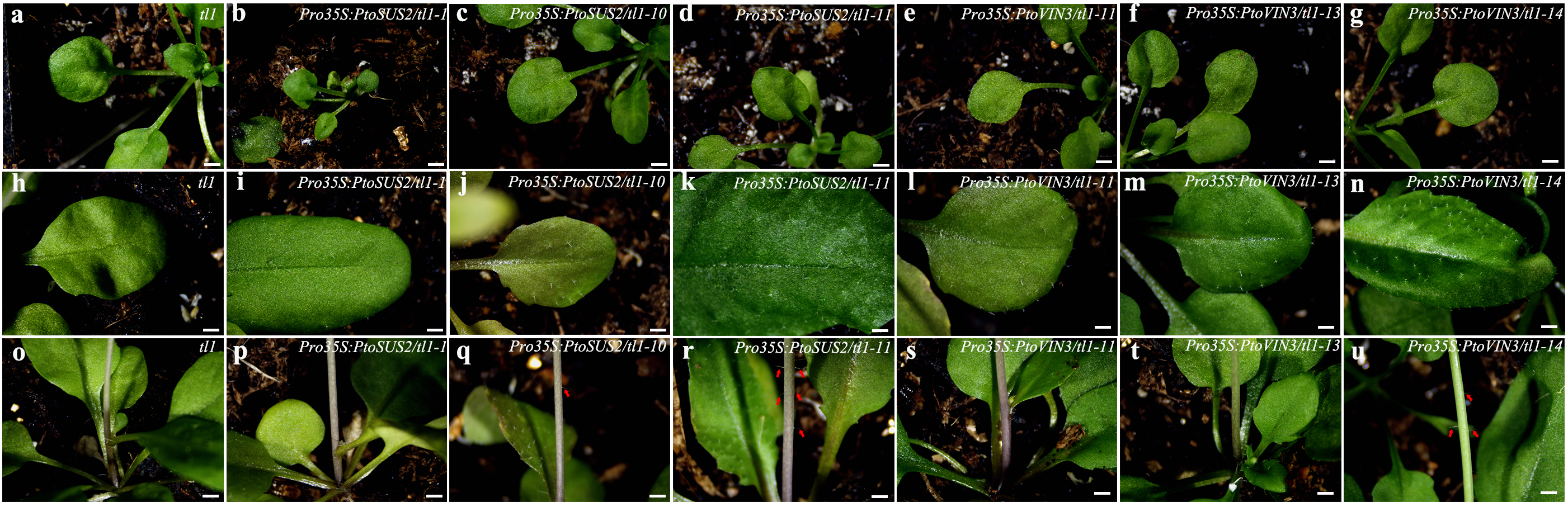

### Figure S9.tif

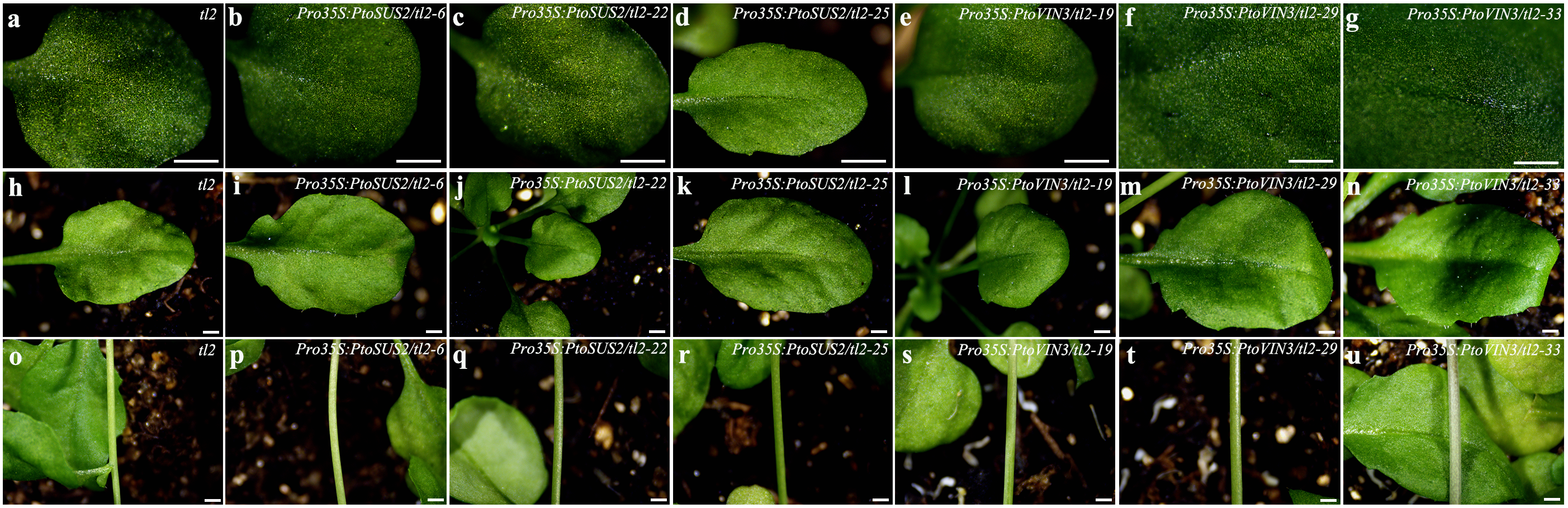

### Figure S10.tif

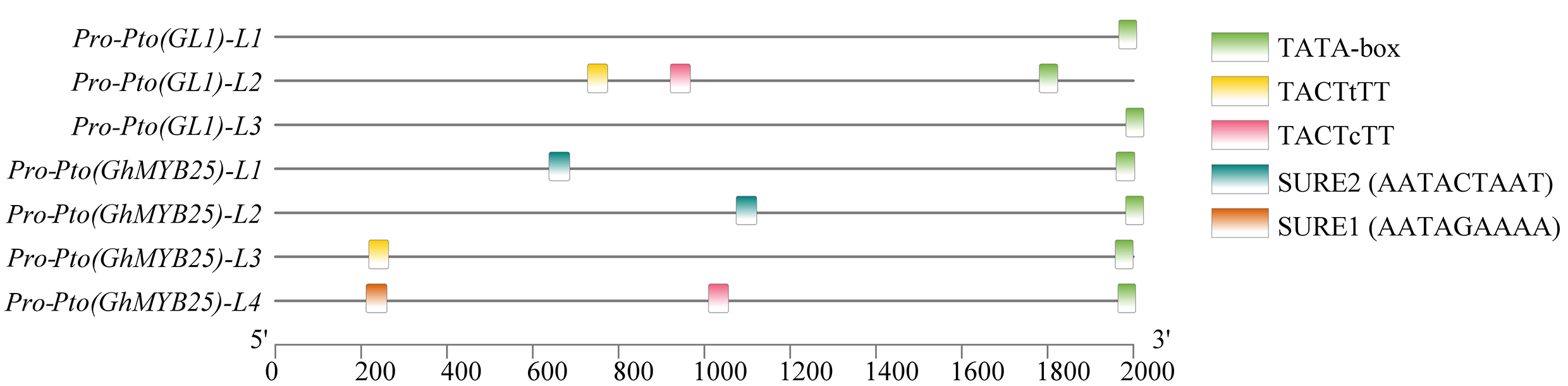
