## Supplementary material for "Sucrose metabolism and candidate genes during catkin fibers development in poplar": Methods S1: Methods S1.docx

**Methods S1: Quantitation of sugar content and enzyme activities**

Samples of 1 g (dry weight, DW) were added to 5 mL distilled water or 90% alcohol, and the mixture was extracted through ultrasonication for 10 min. Subsequently the mixture was centrifuged at 4℃ and 12,000 rpm for 10 min. The mixture was filtered through a 0.45-μm hydrophilic filter prior to analysis. The concentrations of sucrose, fructose and glucose were analyzed by high-performance liquid chromatography (HPLC) using an Agilent 1260 HPLC system (Agilent Technologies, Santa Clara, CA, USA), and a carbohydrate column (Agilent Technologies) was maintained at 35 ℃. The samples were eluted with acetonitrile: distilled water at an 80:20 ratio. The injection volume was 10 μL, and the flow rate was 1 mL min ^−1^. The sugar concentrations were quantified according to a standard solution (Sigma, St. Louis, MO, USA), and defined in terms of mg•g^-1^ DW.

A sample of 5 g (fresh weight, FW) was weighed and ground in liquid nitrogen for 5 min, and then extraction buffer (2 mL volume containing 200 mmol HEPES–NaOH, 0.1% β-mercaptoethanol, 10 mmol sodium ascorbate, 10 mmol cysteine-HCl, 2% glycerol, 0.05% bovine serum albumin [BSA], 1 mmol ethylenediaminetetraacetic acid [EDTA], 1 mmol ethylene glycol-bis(β-aminoethyl ether)-*N,N,N′,N′*-tetraacetic acid [EGTA], 5 mmol MgCl2, 2% polyvinylpolypyrrolidone [PVPP] and 0.05% Triton-X 100, pH 7.5) was added, followed by mixing and centrifuging at 20,000 rpm for 30 min. Subsequently, the supernatant (1 mL) was desalted using a Sephadex G25 PD-10 column with desalination buffer (2 mL volume containing 20 mmol HEPES–NaOH, 0.01% β-mercaptoethanol, 0.2% glycerol, 0.05% BSA, 1 mmol EDTA, 1 mmol EGTA and 0.25 mmol MgCl2, pH 7.5). For SUS analysis, 1 mL of reaction mixture containing 80 mmol 2-ethanesulfonic acid (MES) buffer (pH 5.5), 100 mmol sucrose, 5 mmol NaF, 5 mmol UDP and 100 µL enzyme extract was incubated at 37℃ for 30 min. Subsequently, the mixture was placing in boiling water for 5 min to stop the reaction. The optical density (OD) of the mixture was measured at 540 nm using an ultraviolet spectrophotometer (Cary 5000, Agilent). For SPS activity, 200 μL reaction mixture containing 100 mmol HEPES-NaOH buffer (pH 7.5), 15 mmol MgCl_2_, 1 mmol EDTA, 4 mmol fructose 6-phosphate, 20 mmol glucose 6-phosphate, 16 mmol UDP-glucose, and 100 μL enzyme extract was used. The reaction mixture was incubated AT 30℃ for 30 min. Subsequently, 200 μL NaOH (5 mol/L) was added, and the mixture was placing in boiling water for 10 min to stop the reaction. After cooling, 3.5 mL of anthrone (0.15%, dissolved in 80% H_2_SO_4_) was added, and the mixture was placed in a 40 ℃ water bath for 20 min. The OD of the mixture was measured at 620 nm using the Cary 5000 instrument.
