## Supplementary material for "Sucrose metabolism and candidate genes during catkin fibers development in poplar": Methods S2: Methods S2.docx

**Methods S2: Quantitation of cellulose content**

Cellulose content was analyzed according to the previous research with some modification. Samples were dried at 80 ℃ overnight firstly, and 1 g samples were ground and added into 2 mL 0.1 M phosphoric acid buffer (pH = 7.2). After incubated at 37 ℃ for 30 min, the samples were added into 2 mL 80% (v/v) ethanol and incubated at 80 ℃ for 1 h. Subsequently, the precipitate was suspended using acetone and dried at 60 °C for 16 h. And then the precipitate was added into 2 mL reaction solution (V_acetic acid_: V_water_: V_nitric acid_ = 8: 2: 1) at 100 ℃ for 30 min. The obtained mixture was dried at 60 ℃ overnight. 2mL 72% (v/v) H_2_SO_4_ was added at room temperature for 1 h, and then 10 mL precooled anthrone solution (0.2 g of anthrone in 100 mL H_2_SO_4_) was added and incubated at 100 ℃ for 20 min. The OD of the mixture was measured at 625 nm using the Cary 5000 instrument, and α-cellulose (Sigma, St. Louis, MO, USA) was used as standard.
