## Supplementary material for "Sucrose metabolism and candidate genes during catkin fibers development in poplar": Table S1: Table S1.docx

**Table S1** **Comparison of potential TFs and other genes in poplar and cotton**

| Gene | Homologous in *Populus trichocarpa* | Homologous in cotton | genes ID  (Genebank or Cottongen) | References |
| --- | --- | --- | --- | --- |
| **Transcription factors** | |  |  |  |
| *Pto(GL1)-L1* | Potri.015G075800 | *GhMYB109*  *GaMYB2*  *GaMYB23* | AJ549758  AY626160  DR453866 | (Suo et al., 2003, Wang et al., 2004, Wang et al., 2013) |
| *Pto(GL1)-L2* | Potri.015G075600 |  |  |  |
| *Pto(GL1)-L3* | Potri.012G080400 |  |  |  |
| *Pto(GhMYB25)-LI* | Potri.017G086300 | *GhMYB25*  *GhMYB25-LIKE*  *GhMML3_A12*  *GhMML4_D12* | EU826465  HM598082  KP998102  Gh_D12G1630 | (Machado et al., 2009, Walford et al., 2011, Wan et al., 2016, Wu et al., 2018) |
| *Pto(GhMYB25)-L2* | Potri.010G165700 |  |  |  |
| *Pto(GhMYB25)-L3* | Potri.008G089700 |  |  |  |
| *Pto(GhMYB25)-L4* | Potri.008G089200 |  |  |  |
| *Pto(GhMYB7)-LIKE* | Potri.001G197000 | *GhMYB7/GhMYB9* | AY518319/  AY518320 | (Hsu et al., 2005) |
| *Pto(GL3)-L1* | Potri.003G128000 | *GaDEL65*  *GhDEL65*  *GhbHLH1* | JN997400  AF336280  FJ358540 | (Wang et al., 2013, Shangguan et al., 2016, Meng et al., 2009) |
| *Pto(GL3)-L2* | Potri.001G103600 |  |  |  |
| *Pto(TTG1)-L1* | Potri.012G006100 | *GhTTG1*  *GhTTG3*  *GaTTG1*  *GaTTG3* | AF530908  AF530911  JQ005876  JQ005877 | (Wang et al., 2013, Humphries et al., 2005) |
| *Pto(TTG1)-L2* | Potri.015G002600 |  |  |  |
| *Pto(GhHD1)-L1* | Potri.011G025000 | *GhHD1-A/D*  *GbML1* | JN585951/  JN585952  AY464063 | (Zhang et al., 2010, Walford et al., 2012) |
| *Pto(GhHD1)-L2* | Potri.004G020400 |  |  |  |
| *Pto(GhHox3)-L1* | Potri.015G034100 | *GhHox3* | AY626159 | (Shan et al., 2014) |
| *Pto(GhHox3)-L2* | Potri.012G038500 |  |  |  |
| *Pto(GL2)-L1* | Potri.001G184100 | *GaHox1*  *GaGL2* | AF530913  EU328266 | (Guan et al., 2008, Wang et al., 2013) |
| *Pto(GL2)-L2* | Potri.003G052400 |  |  |  |
| *Pto(GhNAC33)-LIKE* | Potri.001G404400 | *GhNAC33* | Gh_A04G1299 | (Sun et al., 2018) |
| **other genes** |  |  |  |  |
| *PtoAnx-LIKE* | Potri.003G200700 | *AnxGb6* | KC316009 | (Huang et al., 2013) |
|  |  | *GhFAnnxA* | ACJ11719 | (Zhang et al., 2018) |
| *PtoAct1-L1* | Potri.001G309500 | *GhACT1* | AY305723 | (Li et al., 2005) |
| *PtoAct1-L2* | Potri.019G010400 |  |  |  |
| *PtoFIM2-LIKE* | Potri.002G251600 | *GhFIM2* | XP_016696213 | (Zhang et al., 2017a) |
| *PtoRDL1-LIKE* | Potri.018G036700 | *GhRDL-related* | AY072821/  AY641990/  AY641991 | (Xu et al., 2013) |
| *PtoCDPK1-Like* | Potri.008G014700 | *GhCDPK1* | EU723087 | (Zhang et al., 2018) |
| *PtoPOX1-LIKE* | Potri.006G069600 | *GhPOX1* | AJ606075 | (Zhang et al., 2018) |
| *PtoKCH-LIKE* | Potri.005G021100 | *GhKCH1*  *GhKCH2* | AY695833  EF432568 | (Zhang et al., 2018) |
| *PtoCTL-LIKE* | Potri.014G146600 | *GhCTL1*  *GhCTL2* | AY291285  AY291286 | (Zhang et al., 2004) |
| *PtoHUB2-LIKE* | Potri.001G008800 | *GhHUB2-A/D* | XM_016856414/  XM_016843047 | (Feng et al., 2018) |
| *PtoKNL1-LIKE* | Potri.001G112200 | *GhKNL1* | KC200250 | (Feng et al., 2018) |
| *PtoXTH1-L1* | Potri.007G008500 | *GhXTH1* | HM749062 | (Feng et al., 2018) |
| *PtoXTH1-L2* | Potri.001G071000 |  |  |  |
| *PtoXTH1-L3* | Potri.003G159700 |  |  |  |
| *Pto1,3-β-G-L1* | Potri.016G057600 | *Gh1,3-β-G* | CAA92278 | (Feng et al., 2018) |
| *Pto1,3-β-G-L2* | Potri.016G057400 |  |  |  |
| *Pto1,3-β-G-L3* | Potri.006G048100 |  |  |  |
| *Pto1,3-β-G-L4* | Potri.001G255100 |  |  |  |
| *Pto1,3-β-G-L5* | Potri.010G142800 |  |  |  |
| *PtoKOR1-LIKE* | Potri.003G151700 | *GhKOR1* | AY574906 | (Feng et al., 2018) |
| *Pto4CL1-L1* | Potri.019G049500 | *Gh4CL1* | FJ479707 | (Feng et al., 2018) |
| *Pto4CL1-L2* | Potri.006G169600 |  |  |  |
| *Pto4CL1-L3* | Potri.018G094200 |  |  |  |
| *PtoCAD6-L1* | Potri.016G078300 | *GhCAD6* | NM_119958 | (Feng et al., 2018) |
| *PtoCAD6-L2* | Potri.009G062800 |  |  |  |
| *PtoCAD6-L3* | Potri.009G063100 |  |  |  |
| *PtoKIS13A1-LIKE* | Potri.008G221400 | *GhKIS13A1* | KP036626 | (Li et al., 2017) |
| *PtoRac1-L1* | Potri.005G242000 | *GhRac1* | AAD47828 | (Kim and Triplett, 2004) |
| *PtoGA20x-LIKE* | Potri.015G134600 | *GhGA20ox1* | FJ623272 | (Xiao et al., 2010) |
| *PtoDET2-LIKE* | Potri.016G110600 | *GhDET2* | AY141136 | (Luo et al., 2007) |
| *PtoGH3-L1* | Potri.001G298300 | *GhGH3.7-A/D* | Gh_A01G0547/  Gh_D01G0559 | (Yu et al., 2018) |
| *PtoGH3-L5* | Potri.011G129700 |  |  |  |
| *PtoGH3-L2* | Potri.001G410400 | *GhGH3.4-A/D*  *GhGH3.5-A/D* | Gh_A11G0443/  Gh_D11G0514  Gh_A11G1993/  Gh_D11G1989 | (Yu et al., 2018) |
| *PtoGH3-L3* | Potri.007G050300 |  |  |  |
| *PtoGH3-L4* | Potri.009G092900 |  |  |  |
| *PtoPME-L1* | Potri.003G002600 | *GhPME32*  *GhPME31*  *GhPME61*  *GhPME73*  *GhPME38* | CotAD_52896  CotAD_57316  CotAD_40855  CotAD_09740  CotAD_38327 | (Li et al., 2016) |
| *PtoPME-L2* | Potri.003G001900 |  |  |  |
| *PtoPME-L3* | Potri.003G002100 |  |  |  |
| *PtoPME-L4* | Potri.001G162600 |  |  |  |
| *PtoPME-L5* | Potri.001G162500 |  |  |  |
| *PtoPME-L6* | Potri.006G256600 |  |  |  |
| *PtoPME-L7* | Potri.003G002800 |  |  |  |
| *PtoPdBG3-LIKE* | Potri.004G086400 | *GhPdBG3-2A/D* | Gh_A07G1597/  Gh_D07G1793 | (Zhang et al., 2017b) |
| *PtoSWEET-L1* | Potri.012G103200 | *GhSWEET5-A/D* | Gh_A03G0461/  Gh_D03G1078 | (Zhang et al., 2017b) |
| *PtoSWEET-L2* | Potri.015G101600 |  |  |  |
| *PtoSWEET-L3* | Potri.001G060900 | *GhSWEET1-A/D* | Gh_A07G0421/  Gh_D07G0486 | (Zhang et al., 2017b, Sun et al., 2019) |
|  |  | *GhSWEET10-A/D* | Gh_A04G0861/  Gh_D04G1360 |  |
|  |  | *GhSWEET12* | MH660711 |  |
| *PtoKT-LIKE* | Potri.013G083400 | *GhKT1* | GhA04G0968/  GhD04G1511 | (Zhang et al., 2018) |
|  |  | *GhKT2* | GhA12G0124/  GhD12G0141 |  |

Genes ID for *Populus trichocarpa* are searched from *PHYTOZOME* (https://phytozome.jgi.doe.gov/) data base, and gene ID for cotton are searched from *Genebank* (https://www.ncbi.nlm.nih.gov/) or *Cottongen* (https://www.cottongen.org/) data base.

**Reference**

**Feng, H., Li, X., Chen, H., Deng, J., Zhang, C., Liu, J., et al.** 2018. GhHUB2, a ubiquitin ligase, is involved in cotton fiber development via the ubiquitin–26S proteasome pathway. *Journal of experimental botany,* **69,** 5059-5075.

**Guan, X. Y., Li, Q. J., Shan, C. M., Wang, S., Mao, Y. B., Wang, L. J., et al.** 2008. The HD‐Zip IV gene GaHOX1 from cotton is a functional homologue of the Arabidopsis GLABRA2. *Physiologia Plantarum,* **134,** 174-182.

**Hsu, C.-Y., Jenkins, J. N., Saha, S. & Ma, D.-P.** 2005. Transcriptional regulation of the lipid transfer protein gene LTP3 in cotton fibers by a novel MYB protein. *Plant Science,* **168,** 167-181.

**Huang, Y., Wang, J., Zhang, L. & Zuo, K.** 2013. A cotton annexin protein AnxGb6 regulates fiber elongation through its interaction with actin 1. *PLos one,* **8**.

**Humphries, J. A., Walker, A. R., Timmis, J. N. & Orford, S. J.** 2005. Two WD-repeat genes from cotton are functional homologues of the Arabidopsis thalianaTRANSPARENT TESTA GLABRA1 (TTG1) gene. *Plant molecular biology,* **57,** 67-81.

**Kim, H. J. & Triplett, B. A.** 2004. Characterization of GhRac1 GTPase expressed in developing cotton (Gossypium hirsutum L.) fibers. *Biochimica et Biophysica Acta (BBA)-Gene Structure and Expression,* **1679,** 214-221.

**Li, W., Shang, H., Ge, Q., Zou, C., Cai, J., Wang, D., et al.** 2016. Genome-wide identification, phylogeny, and expression analysis of pectin methylesterases reveal their major role in cotton fiber development. *BMC genomics,* **17,** 1000.

**Li, X.-B., Fan, X.-P., Wang, X.-L., Cai, L. & Yang, W.-C.** 2005. The cotton ACTIN1 gene is functionally expressed in fibers and participates in fiber elongation. *The Plant Cell,* **17,** 859-875.

**Li, Y.-J., Zhu, S.-H., Zhang, X.-Y., Liu, Y.-C., Xue, F., Zhao, L.-J., et al.** 2017. Expression and functional analyses of a Kinesin gene GhKIS13A1 from cotton (Gossypium hirsutum) fiber. *BMC biotechnology,* **17,** 50.

**Luo, M., Xiao, Y., Li, X., Lu, X., Deng, W., Li, D., et al.** 2007. GhDET2, a steroid 5α‐reductase, plays an important role in cotton fiber cell initiation and elongation. *The Plant Journal,* **51,** 419-430.

**Machado, A., Wu, Y., Yang, Y., Llewellyn, D. J. & Dennis, E. S.** 2009. The MYB transcription factor GhMYB25 regulates early fibre and trichome development. *The Plant Journal,* **59,** 52-62.

**Meng, C.-M., Zhang, T.-Z. & Guo, W.-Z.** 2009. Molecular cloning and characterization of a novel Gossypium hirsutum L. bHLH gene in response to ABA and drought stresses. *Plant molecular biology reporter,* **27,** 381-387.

**Shan, C.-M., Shangguan, X.-X., Zhao, B., Zhang, X.-F., Chao, L.-M., Yang, C.-Q., et al.** 2014. Control of cotton fibre elongation by a homeodomain transcription factor GhHOX3. *Nature communications,* **5,** 5519.

**Shangguan, X. X., Yang, C. Q., Zhang, X. F. & Wang, L. J.** 2016. Functional characterization of a basic Helix‐Loop‐Helix (bHLH) transcription factor GhDEL65 from cotton (Gossypium hirsutum). *Physiologia plantarum,* **158,** 200-212.

**Sun, H., Hu, M., Li, J., Chen, L., Li, M., Zhang, S., et al.** 2018. Comprehensive analysis of NAC transcription factors uncovers their roles during fiber development and stress response in cotton. *BMC plant biology,* **18,** 150.

**Sun, W., Gao, Z., Wang, J., Huang, Y., Chen, Y., Li, J., et al.** 2019. Cotton fiber elongation requires the transcription factor Gh MYB 212 to regulate sucrose transportation into expanding fibers. *New Phytologist,* **222,** 864-881.

**Suo, J., Liang, X., Pu, L., Zhang, Y. & Xue, Y.** 2003. Identification of GhMYB109 encoding a R2R3 MYB transcription factor that expressed specifically in fiber initials and elongating fibers of cotton (Gossypium hirsutum L.). *Biochimica et Biophysica Acta (BBA)-Gene Structure and Expression,* **1630,** 25-34.

**Walford, S. A., Wu, Y., Llewellyn, D. J. & Dennis, E. S.** 2011. GhMYB25‐like: a key factor in early cotton fibre development. *The Plant Journal,* **65,** 785-797.

**Walford, S. A., Wu, Y., Llewellyn, D. J. & Dennis, E. S.** 2012. Epidermal cell differentiation in cotton mediated by the homeodomain leucine zipper gene, GhHD‐1. *The Plant Journal,* **71,** 464-478.

**Wan, Q., Guan, X., Yang, N., Wu, H., Pan, M., Liu, B., et al.** 2016. Small interfering RNA s from bidirectional transcripts of Gh MML 3_A12 regulate cotton fiber development. *New Phytologist,* **210,** 1298-1310.

**Wang, G., Zhao, G. H., Jia, Y. H. & Du, X. M.** 2013. Identification and Characterization of Cotton Genes Involved in Fuzz‐F iber Development. *Journal of integrative plant biology,* **55,** 619-630.

**Wang, S., Wang, J.-W., Yu, N., Li, C.-H., Luo, B., Gou, J.-Y., et al.** 2004. Control of plant trichome development by a cotton fiber MYB gene. *The Plant Cell,* **16,** 2323-2334.

**Wu, H., Tian, Y., Wan, Q., Fang, L., Guan, X., Chen, J., et al.** 2018. Genetics and evolution of MIXTA genes regulating cotton lint fiber development. *new phytologist,* **217,** 883-895.

**Xiao, Y.-H., Li, D.-M., Yin, M.-H., Li, X.-B., Zhang, M., Wang, Y.-J., et al.** 2010. Gibberellin 20-oxidase promotes initiation and elongation of cotton fibers by regulating gibberellin synthesis. *Journal of plant physiology,* **167,** 829-837.

**Xu, B., Gou, J.-Y., Li, F.-G., Shangguan, X.-X., Zhao, B., Yang, C.-Q., et al.** 2013. A cotton BURP domain protein interacts with α-expansin and their co-expression promotes plant growth and fruit production. *Molecular Plant,* **6,** 945-958.

**Yu, D., Qanmber, G., Lu, L., Wang, L., Li, J., Yang, Z., et al.** 2018. Genome-wide analysis of cotton GH3 subfamily II reveals functional divergence in fiber development, hormone response and plant architecture. *BMC plant biology,* **18,** 1-18.

**Zhang, D., Hrmova, M., Wan, C.-H., Wu, C., Balzen, J., Cai, W., et al.** 2004. Members of a new group of chitinase-like genes are expressed preferentially in cotton cells with secondary walls. *Plant molecular biology,* **54,** 353-372.

**Zhang, F., Jin, X., Wang, L., Li, S., Wu, S., Cheng, C., et al.** 2018. A Cotton Annexin Affects Fiber Elongation and Secondary Cell Wall Biosynthesis Associated with Ca2+ Influx, ROS Homeostasis, and Actin Filament Reorganization. The authors have chosen to retract the above article from Plant Physiology follow-ing an investigation into concerns centered on the origins and assembly of.

**Zhang, F., Zuo, K., Zhang, J., Liu, X., Zhang, L., Sun, X., et al.** 2010. An L1 box binding protein, GbML1, interacts with GbMYB25 to control cotton fibre development. *Journal of experimental botany,* **61,** 3599-3613.

**Zhang, M., Han, L. B., Wang, W. Y., Wu, S. J., Jiao, G. L., Su, L., et al.** 2017a. Overexpression of GhFIM2 propels cotton fiber development by enhancing actin bundle formation. *Journal of integrative plant biology,* **59,** 531-534.

**Zhang, Z., Ruan, Y.-L., Zhou, N., Wang, F., Guan, X., Fang, L., et al.** 2017b. Suppressing a putative sterol carrier gene reduces plasmodesmal permeability and activates sucrose transporter genes during cotton fiber elongation. *The Plant Cell,* **29,** 2027-2046.
