## Supplementary material for "Sucrose metabolism and candidate genes during catkin fibers development in poplar": Table S2: Table S2.docx

**Table S2 Primers used for RT-qPCR**

| **Gene** | **Forward primer (5’→3’)** | **Reverse primer (5’→3’)** |
| --- | --- | --- |
| Primers used for poplar | | |
| *PtActin* | GGATGGCTATATAGGTTGCAGGAA | GAGATGCTGGGAAGGAATTGAAAG |
| *PtoCesA1/PtoCesA11* | CGTACCTCCTTCCTCGAGTCCA | AGCCCTTAAATGCAGCTAACTTT |
| *PtoCesA10* | GAGCTCAGATGGAGCGGAG | CTCTCCAAAATCGTCATCGTCT |
| *PtoCesA2* | CCATGTTTCTGGGTTCGCT | AGGCATTGAAGAATCAGAAAACG |
| *PtoCesA5* | GGTAATTGTCCACCTTTACCCG | CACAATTCAACCCACAAATTTCT |
| *PtoCesA6* | CGAGGAGGATGCCGAGATTTC | CTGTCCTTCCATGCCACACTC |
| *PtoCesA9* | TGTCAGCACAAACTTCACTGTG | GATGATGACCCACAAGGCG |
| *PtoCesA15* | GGTCTTGGAGGACCTGAATCC | TTCCACTGCCACCAATAGGAG |
| *PtoCesA16* | CAGCACCATGGTGGACCTG | TTCCCGTCACCACCGACC |
| *PtoCesA4* | TCGATGGAGTGGAGTCAGCAT | GTACAGCTCTCCAAACTCGGC |
| *PtoCesA8* | CCAACTCCTCAACCGCCATG | AGCTAGGGTGTCTTGGTTGTCT |
| *PtoCesA18* | CACCGACACACCCTAGCTTTG | AGAGGTGCGTAACAAACAAATGA |
| *PtoCesA7* | CGTCCCCATTTTCAATATGGAG | AAGAGTTGCAGGATTGGTTGTC |
| *PtoCesA17* | TGCCATTTGCCTAGTCACTGG | CTGAGGTGCCACCGATCAC |
| *PtoSPS1A* | GGCTGTATATGTCAACCTGGGA | AGGATGGTCATCATTTCCTTC |
| *PtoSPS1B* | GGAAATGAAGACCATCCCTCTT | CCGACATTCTCCAAATGCTG |
| *PtoSPS2* | GGATCAAACGAGGAGTGAGCTG | AAATTGGTGGATCAGGAGACGC |
| *PtoSPS3A* | CCCTGAAGTTGATTGGAGCTAC | GGCCACAGTAATTCTTTTTGGAG |
| *PtoSPS3B* | CTAACAGGACACTCACTCGGGC | ACGCAAGACTCTTTCAAGCTTG |
| *PtoSPP1* | GGAACCTAGTCACTGAGGAAACA | ATAGCGCCTTCGTCACATTC |
| *PtoSPP2* | ATCCATCTGGTGCTGAACTTCC | ACCAACCATGTATCCAGGCCA |
| *PtoSPP3* | TCTGCTAGTTTTCTCCACCGGA | TGGGCTGTTTCCTCTGTGACTA |
| *PtoSUS1* | CATCGATCCTTATCATGGCGAG | TGTACCGACGGCTCTCAAGAC |
| *PtoSUS2* | CACTGAGGAGAAACTTAGGCTGA | CAAGATTAGCTAACTCGCGCAG |
| *PtoSUS3* | GGTATGTGGAGCAGGGAAAAGG | TCCCAACTGAAGGTGGAGGAAT |
| *PtoSUS4* | GCTTCGGTTGGCACTGATAG | TCCCTTCTCGAAGCCCCACA |
| *PtoSUS5* | AAGCAACAAGGCCTAAATGTG | CGGAGGACCCTATTCTCTACCC |
| *PtoSUS6* | TGGGGTCTCAGGATTCCACATT | GCCTCCAAAACCCGTAAACAGA |
| *PtoSUS7* | ACAAGGAACAGAAGCTTGCGAA | TGGGTTGGAAGTGAATGCTGAG |
| *PtCWIN3* | CAAGACATGAACAAAGGATGGG | AGTTGAACATTGTGGCTCCTCAG |
| *PtCWIN4* | CCTCCTTTCACTTCCAACCACC | TGCGCCCATATCATAAAATCACCA |
| *PtCWIN5* | AACAGGCGAATTTTGTGGGGAT | CCGGCCACTGTACTAATTGCTT |
| *PtVIN2* | AACAATGGAGCTCGACATGTCC | TCTTTCACCTGAGCTCCCTTCA |
| *PtVIN3* | CCTCCTCTGACCTTGAACATGC | GCGTTGAAGATGACCGGTTTTG |
| *PtoNIN2* | TGGAGTCTGAGGATTGGCGT | TAAAGTCGGGATTGCTTTCCA |
| *PtoNIN5* | ATGTCTCTTTTTGGCCAATCTC | AAGGCTTAATACCGATCCCATT |
| *PtoNIN1* | TGATTCTGGGCTGTGGTGGATA | CTGGACATCAACCCTCTCTTGC |
| *PtoNIN1* | CAGTTGCAGAGCTGGGAGAAAA | TATCCACCACAGCCCAGAATCA |
| *PtoNIN3* | ATGGCGACTTCGGATGCAGT | ACAGATCACCGTCACCTGATCT |
| *PtoNIN4* | CGTTTGGATGGTGATGATGATTT | GCACATCAATCCTCTCCTGAAG |
| *PtoNIN9* | TGAAGAGTGTCGATGCCCATC | GAAAGCAAAGGCACCCCAA |
| *PtoNIN11* | AAGCTTGAGAAATGCTGATACCC | GAAGGAATTGGCAACCCAGTAG |
| *PtoNIN12* | GATGGGGGGGCTTAGGAAC | TCTTGCAAGCCCGATCGAG |
| *PtoNIN8* | GAAACGGTGGGGCTTATGAAT | CAATTCCTTTTGCAAGCCCA |
| *Pto(GL1)-L1* | AAGTGGAATCGCCTCAAGAGAA | GGGACCCGACCAGCAATTAAG |
| *Pto(GL1)-L2* | GGTCTCTAATTGCTGGTCGGATT | CGCCTCCATTTTTTTGGAGGA |
| *Pto(GL1)-L3* | GTCTTTAATTGCCGGTCGAGTT | CGCCCCTAATTCTTCAGAAAA |
| *Pto(GhMYB25)-LI* | ACTCCATGCCCTCTTAGGGAAC | GGATTGACCGTCAATGGAAAGT |
| *Pto(GhMYB25)-L2* | ATGCGTGGTTTGACTCTGCAAC | TTCCCGTTCTCTTCAAGATTCC |
| *Pto(GhMYB25)-L3* | AAACGCACCTATGGGGAATATC | CAGATGACCCAGTTGGAGAGG |
| *Pto(GhMYB25)-L4* | TTGAGAAACTCAAAGAACCAGCAC | ATGATAGGTGCGTTTTCATTCG |
| *Pto(GhMYB7)-LIKE* | GTCCACAATTCGAGCGTTCCT | TTGAGGTTGGTGAAGGGAGGT |
| *Pto(GL3)-L1* | CTAGACACTGTTTCCCCTGATGA | GATGATGAACACCGTTGCTGAA |
| *Pto(GL3)-L2* | AGACATAACTTCCCCTAACCGC | GGAATGATGGACACTGTGGTC |
| *Pto(TTG1)-L1* | GCTGAGTTGGAGAGGCATATGG | TCAGAAGTAGCGGAGCACATAGA |
| *Pto(TTG1)-L2* | GGAGTTGGAGAGGCATCGTT | CAGAAGCAGCAGAGCACATG |
| *Pto(GhHD1)-L1* | ATGTTCCACGCAAATATGTTTGA | GTGTTGTTCACTGGGATCTTGG |
| *Pto(GhHD1)-L2* | TAGCCACCATATGCTTGATATGAC | TGGCCGCTGGGATCCTGA |
| *Pto(GhHox3)-L1* | CTAGTGGCGGTGGTGGTGGA | CATCCACAAAGGTTCGTTGGT |
| *Pto(GhHox3)-L2* | TCAAGAGAGCTGCATCGACTC | ACCATCTCCTTGGTCAGGATAA |
| *Pto(GL2)-L1* | AAATCAAGGCACGTACGTTGG | CTGTTGTTCCTCAGTGCTTGTCA |
| *Pto(GL2)-L2* | AGAGACCAGGGTTGTGTTTGCG | CACTGCCCCATTTCTATTTGCA |
| *Pto(GhNAC33)-LIKE* | TGTGCTTGGATCAATACCGCC | CGAAGAGGCCATGTGAGACAA |
| *PtoAnx-LIKE* | TGATCATGGCTCTTCCATCAC | TCCTCATCTGTCCCAACCTTT |
| *PtoAct1-L1* | CGAATGAGCAAGGAGATCACC | GCTGAGGGAAGCAAGGATAGAT |
| *PtoAct1-L2* | CTTACAGAGGCTCCTCTTAACCCT | GACACACCATCACCAGAATCCAG |
| *PtoFIM2-LIKE* | TCGGAGAAGGCATCCTATGTT | TTCGCAATCTCGAAGAGATCA |
| *PtoRDL1-LIKE* | GCCTTGCCACCTGAACTTTAC | GTTCCTCCAGGCTTTCCTTTT |
| *PtoCDPK1-Like* | GTAGGACCGAGCATTTCCAAG | GGTTTTGGGATGGTCATCTGT |
| *PtoPOX1-LIKE* | GTGCAAAGCCAATTGAAAACA | ACCCCGCAATCAATACTGAAC |
| *PtoKCH-LIKE* | GCAACAAATTTGGCCTTTCTC | ACTCCTGCTGTTTTTCGAAGC |
| *PtoCTL-LIKE* | GGATGGGAGTGCTTAGACTGG | CGGTGGTACAAAACCCAAGAT |
| *PtoHUB2-LIKE* | ACGGTGGGAATTGATGGATGC | ACCTCCATGAGCTCTTCCTCC |
| *PtoKNL1-LIKE* | AAGTGGCCTTACCCAACTGAA | TTGAGAGTTGCTGTGCCAGTT |
| *PtoXTH1-L1* | TACTTGTTCGGCCGTGTTAGC | TGCAGCAGGATCAAACCACAG |
| *PtoXTH1-L2* | ACCCAACCATAAGATACCACTCT | GGCTGGTTGAAAGGAAATTTC |
| *PtoXTH1-L3* | ACATGGCTGCTGCTTATCCG | GAACATAGTTCCTACCGAACGC |
| *Pto1,3-β-G-L1* | TGGTAACCTACCACCGGCACA | GTTGCCAAAACTCCTTACATTTC |
| *Pto1,3-β-G-L2* | GGTAACTTGCCACCACCATC | TTGCCAAATTGTTTTACATTGC |
| *Pto1,3-β-G-L3* | GCATCACATCCAGCTGGCAA | CATTCTTTGAATGCCTCTCTCGT |
| *Pto1,3-β-G-L4* | AGAGACACCCTTGAAGCCCTT | GCAATGTAGCGGAACCTGACA |
| *Pto1,3-β-G-L5* | TTCCTTAACGGCTGCAATGCT | TGCCTGAACCTCTGAGAGCAT |
| *PtoKOR1-LIKE* | GAACAAAAGGAGGCTTGATCG | TTTGGTCCACAATACCATCCA |
| *Pto4CL1-L1* | TCCGTGGCCACGGTTGAG | GGAGGTGGTTCGAGATGGTT |
| *Pto4CL1-L2* | CGCAAACCCCTTCTACACTCCC | CTGGTGGAGAGTCAACGGTCAC |
| *Pto4CL1-L3* | TTTTGCATTTCTTGGAGCGTCGT | CATGGTCGTTTTCTTGAGCAAAC |
| *PtoCAD6-L1* | GGCTAGGGATCAATCTGGTCAT | AGCCCCAGTCATTCTTGATACTG |
| *PtoCAD6-L2* | GCACAATTAGCGTGTACATTG | GGTAAGACGCGGACATCCC |
| *PtoCAD6-L3* | CCAGTGTTTCCTCTGATCATGGG | TTCACATAATCCATCGAGATAACC |
| *PtoKIS13A1-LIKE* | CTAGTCAGCAAGCGGGGAATG | ATGAACATGAATCCCCACCGC |
| *PtoRac1-L1* | TATCGAGGGGCAGATGTCTTT | TATTAATCGGCACTGCACCAG |
| *PtoGA20x-LIKE* | TCGCTTTCTTTCTGTGTCCAA | GAGCAACATAGGCCATGTGAA |
| *PtoDET2-LIKE* | CCCCTACATTCATTTCCCTGA | GAATGAGAGCTTTGGGATTGG |
| *PtoGH3-L1* | TGCCAATGGTGATACTTCACAG | GGAACAAACTGAGTCATAACAGGC |
| *PtoGH3-L2* | GAAATAACTGACCCTTCCGTCA | TGCACGCCAAAGGTAAACC |
| *PtoGH3-L3* | AGCAATGCCGAGACTGAATATC | AAGATGGGAGACTTGTCACCAT |
| *PtoGH3-L4* | TGCCGACTCCGTTCAAGACA | GATCCTCGTATCTAATCATGGGG |
| *PtoGH3-L5* | CAACGGTGATGCCTCGCCA | GGCCAGGAACAAACTGGCTCATT |
| *PtoPME-L1* | TTGAGGGGATTCAGGGATATT | GATGATTGGAACAGCCGTCT |
| *PtoPME-L2* | CATGGCTTTCTAGGCAAGTAAG | GCTGGAGCGTCTTTGATCAAC |
| *PtoPME-L3* | CATCGTTTTTGAGGGGATTCA | TTTAGTGCCTTCGGTCAACGT |
| *PtoPME-L4* | CACTCACCAATCAGCAAACTG | CCATCATCTTGATCGCTAAGAG |
| *PtoPME-L5* | AACTCACCAATCAGCAAAATG | GCATCTTGATCATCGCTAAGAG |
| *PtoPME-L6* | GCTGATCATGCCAATCTCGTC | ATTGATAGCAGTCGTTTGTCGA |
| *PtoPME-L7* | GAGATGGGAAACTGCGGG | AGCTGCCACCACCACGTT |
| *PtoPdBG3-LIKE* | TTTTCACAGCTGTTCCCAATG | TGAGCATGGGAATCAAAACAG |
| *PtoSWEET-L1* | ACTGGGCTTTATCCTCGGTCT | CATGATTTCCAGGGCCTGTCC |
| *PtoSWEET-L2* | GAGATGGCCTTGCACTTGACA | GTGGCTGGTCTTCTTCTTGCA |
| *PtoSWEET-L3* | GCAACAAGAGAGAGCAGGGTC | GCCTTGACACGAGTTGAACCA |
| *PtoKT-LIKE* | CCTGGCATTGGCTTAGTTTTC | AAAAGGCACAGGCACTGATTT |
| Primers used for transgenic plants | | |
| *AtMYB23* | TCCGGGAAGAACAGACAACCAAGT | AGGACGTTGTGGTTATAAGGGCCA |
| *AtGL2* | AGGCTATTCAAGAACGGCACGAGA | AGCTTATCGAGCTCGGCTTTCAGT |
| *AtGL3* | ACGGCACCATCACAAGGAATGT | TTCAATTTCTCGCGGCGCTT |
| *AtTTG1* | CTCCGCTCATCCTCCGGTCACAG | CAACGGCGCACAAAACTCGCTCG |
| *AtGL1* | GTGAACAAAGGCAATTTCACTG | GTTCTTCCCGGTACTCTTTTAGC |
| *AtEF1* | TTGACCAGATCAACGAGCCCAAGA | ACTCGTGGTGCATCTCAACAGACT |
| *N-PtoSUS2* | GAACTTGTTGATGGAGGCTCC | CCTTGTGACAGTGCACCTTG |
| *N-PtoVIN3* | CTTCAACGCCAAGACCTACAGAG | AGACCTTATCGCTGAAAAATGATG |
