## Supplementary material for "Sucrose metabolism and candidate genes during catkin fibers development in poplar": Table S3: Table S3.docx

**Table S3 Accession numbers of genes used for phylogenetic tree construction**

| **Gene** | **Accession number** | **Gene** | **Accession number** |
| --- | --- | --- | --- |
| **Acid invertase** | | *PtrSUS2* | Potri.006G136700.1 |
| *AtVI1* | AT1G62660.1 | *PtrSUS3* | Potri.002G202300.1 |
| *AtVI2* | AT1G12240.1 | *PtrSUS4* | Potri.017G139100.1 |
| *AtCWINV1* | AT3G13790.1 | *PtrSUS5* | Potri.004G081300.1 |
| *AtCWINV2* | AT3G52600.1 | *PtrSUS6* | Potri.012G037200.1 |
| *AtCWINV3* | AT1G55120.1 | *PtrSUS7* | Potri.015G029100.1 |
| *AtCWINV4* | AT2G36190.1 | **Cellulose synthase** | |
| *AtCWINV5* | AT3G13784.1 | *AtCesA1* | AT4G32410.1 |
| *GhCWIN1* | AI725433 | *AtCesA2* | AT4G39350.1 |
| *GhVIN1* | FJ915120 | *AtCesA3* | AT5G05170.1 |
| *GhVIN2* | GU252170 | *AtCesA4* | AT5G44030.1 |
| *GhVIN3* | FJ864677 | *AtCesA5* | AT5G09870.1 |
| *PtrCWIN1* | Potri.016G077400.1 | *AtCesA6* | AT5G64740.1 |
| *PtrCWIN2* | Potri.016G077500.1 | *AtCesA7* | AT5G17420.1 |
| *PtrCWIN3* | Potri.006G210600.1 | *AtCesA8* | AT4G18780.1 |
| *PtrCWIN4* | Potri.006G227500.1 | *AtCesA9* | AT2G21770.1 |
| *PtrCWIN5* | Potri.006G227400.1 | *AtCesA10* | AT2G25540.1 |
| *PtrVIN1* | Potri.003G126300.1 | *GhCesA1* | U58283 |
| *PtrVIN2* | Potri.003G112600.1 | *GhCesA2* | U58284 |
| *PtrVIN3* | Potri.015G127100.1 | *GhCesA2-A_T_* | JN382209 |
| **Alkaline or neutral invertase** | | *GhCesA2-D_T_* | JN382210 |
| *AtCINV1* | AT1G35580.1 | *GhCesA3* | AF150630 |
| *AtCINV2* | AT4G09510.1 | *GhCesA4* | AF413210 |
| *AtINVA* | AT1G56560.1 | *GhCesA5* | JQ345693 |
| *AtINVB* | AT4G34860.1 | *GhCesA6* | JQ345694 |
| *AtINVC* | AT3G06500.1 | *GhCesA7* | JQ345695 |
| *AtINVD* | AT1G22650.1 | *GhCesA8* | JQ345696 |
| *AtINVE* | AT5G22510.1 | *GhCesA9* | JQ345699 |
| *AtINVF* | AT1G72000.1 | *GhCesA10* | JQ345697 |
| *AtINVH* | AT3G05820.1 | *PtCesA1* | AF072131 |
| *PtrNIN1* | Potri.008G101500.1 | *PtCesA2* | AY095297 |
| *PtrNIN2* | Potri.013G006600.1 | *PtCesA3* | AF527387 |
| *PtrNIN3* | Potri.008G024100.1 | *PtCesA4* | AY162181 |
| *PtrNIN4* | Potri.010G236100.1 | *PtCesA5* | AY055724 |
| *PtrNIN5* | Potri.005G010800.1 | *PtCesA6* | AY196961 |
| *PtrNIN6* | Potri.004G186500.1 | *PtCesA7* | AY162180 |
| *PtrNIN7* | Potri.005G239400.1 | *PttCesA1* | AY573571 |
| *PtrNIN8* | Potri.019G082000.1 | *PttCesA2* | AY573572 |
| *PtrNIN9* | Potri.004G167500.1 | *PttCesA3-1* | AY573573 |
| *PtrNIN10* | Potri.002G173600.1 | *PttCesA3-2* | AY573574 |
| *PtrNIN11* | Potri.009G129000.1 | *PttCesA4* | AY573575 |
| *PtrNIN12* | Potri.013G110800.1 | *PtrCesA2* | Potri.002G066600.1 |
| **Sucrose synthase** | | *PtrCesA3* | Potri.001G266400.1 |
| *AtSUS1* | AT5G20830.1 | *PtrCesA4* | Potri.002G257900.1 |
| *AtSUS2* | AT5G49190.1 | *PtrCesA5* | Potri.005G194200.1 |
| *AtSUS3* | AT4G02280.1 | *PtrCesA6* | Potri.005G087500.1 |
| *AtSUS4* | AT3G43190.1 | *PtrCesA7* | Potri.006G251900.1 |
| *AtSUS5* | AT5G37180.1 | *PtrCesA8* | Potri.011G069600.1 |
| *AtSUS6* | AT1G73370.1 | *PtrCesA9* | Potri.007G076500.1 |
| *GaSUS1* | JQ995522 | *PtrCesA10* | Potri.018G103900.1 |
| *GaSUS2* | JQ995523 | *PtrCesA1/PtrCesA11* | Potri.006G181900.1 |
| *GaSUS3* | JQ995524 | *PtrCesA12* | Potri.006G052600.1 |
| *GaSUS4* | JQ995525 | *PtrCesA13* | Potri.009G060800.1 |
| *GaSUS5* | JQ995526 | *PtrCesA14* | Potri.016G054900.1 |
| *GaSUS6* | JQ995527 | *PtrCesA15* | Potri.013G019800.1 |
| *GaSUS7* | JQ995528 | *PtrCesA16* | Potri.005G027600.1 |
| *GhSUS3* | U73588 | *PtrCesA17* | Potri.018G029400.1 |
| *PtrSUS1* | Potri.018G063500.1 | *PtrCesA18* | Potri.004G059600.1 |

The accession numbers in Arabidopsis were searched in *TAIR* (https://www.arabidopsis.org/), the accession numbers in *Populus trichocarpa* were searched in *PHYTOZOME V12* (https://phytozome.jgi.doe.gov/), and the accession numbers in cotton and other poplars were searched in *Genebank* (<https://www.ncbi.nlm.nih.gov/>). At, *Arabidopsis thaliana*; Gh, *Gossypium hirsutum*; Pto, *Populus tomentosa*; Ptr, *Populus trichocarpa*; Ptt, *Populus tremula × Populus tremuloides*; Pt, *Populus tremuloides*.
