## Supplementary material for "Sucrose metabolism and candidate genes during catkin fibers development in poplar": Table S4: Table S4 .docx

**Table S4** **Comparison of the TFs for YIH in poplar and Arabidopsis**

| Gene | Homologous in *Populus trichocarpa* | Homologous in *Arabidopsis thaliana* | genes ID  (TAIR) |
| --- | --- | --- | --- |
| *Pto(GL1)-L3* | Potri.012G080400 | *AtGL1* | AT3G27920 |
| *Pto(GhMYB25)-L2* | Potri.010G165700 |  |  |
| *Pto(GhMYB25)-L4* | Potri.008G089200 |  |  |
| *Pto(GL3)-L1* | Potri.003G128000 | *AtGL3* | AT5G41315 |
| *Pto(DRMY1)-LIKE* | Potri.002G106500 | *AtDRMY1* | AT1G58220 |
| *Pto(ABS5)-LIKE* | Potri.013G117600 | *AtABS5* | AT1G68810 |
| *Pto(MYC2)-LIKE* | Potri.001G142200 | *AtMYC2* | AT1G32640 |
| *Pto(MYB109)-LIKE* | Potri.008G062700 | *AtMYB109* | AT3G55730 |

Genes ID for *Populus trichocarpa* are searched from *PHYTOZOME V12* (https://phytozome.jgi.doe.gov/) data base, and gene ID for *Arabidopsis thaliana* are searched from *TAIR* (https://www.arabidopsis.org/) data base.
