## Supplementary material for "Sucrose metabolism and candidate genes during catkin fibers development in poplar": Table S5: Table S5.docx

**Table S5 Primers used for plasmid construction**

| **Gene** | **Forward primer (5’→3’)** | **Reverse primer (5’→3’)** |
| --- | --- | --- |
| Primers used for overexpression plasmid | | |
| *PtoSUS2* | AGAACACGGGGGACTCTTGACATGGCTGCACTTACTCGTGTCC | GGGGAAATTCGAGCTGGTCACTTACTCGATAGTCAAAGGAACAGA |
| *PtoVIN3* | AGAACACGGGGGACTCTTGACATGGAGACTAACCCCTCCCATA | GGGGAAATTCGAGCTGGTCACTCACATTGATGTTATCACTCTTCTT |
| Primers used for Y1H plasmid | | |
| *Pto(GL1)-L3* | GCCATGGAGGCCAGTGAATTCATGCAAGGAGCTGGGAAT | CAGCTCGAGCTCGATGGATCCTTACAAGTTATACCAAAAAAG |
| *Pto(GhMYB25)-L2* | GCCATGGAGGCCAGTGAATTCATGGTAAGGTCTCAGTGC | CAGCTCGAGCTCGATGGATCCTCAAAACACAGATGACGT |
| *Pto(GhMYB25)-L4* | GCCATGGAGGCCAGTGAATTCATGGTTAAGTCTCAATGC | CAGCTCGAGCTCGATGGATCCTCAAAATACAGATGACCC |
| *Pto(GL3)-L1* | GCCATGGAGGCCAGTGAATTCATGGAAAATGAGCTACGT | CAGCTCGAGCTCGATGGATCCTCATCGATTCGAGAACAA |
| *Pto(DRMY1)-LIKE* | GCCATGGAGGCCAGTGAATTCATGATTGAGAAATCCAAG | CAGCTCGAGCTCGATGGATCCCTATCCGTCCATCATTCC |
| *Pto(ABS5)-LIKE* | GCCATGGAGGCCAGTGAATTCATGGAACCTTGTTCTTGG | CAGCTCGAGCTCGATGGATCCTCAACTGCAGAGATCCCC |
| *Pto(MYC2)-LIKE* | GCCATGGAGGCCAGTGAATTCATGACTGATTACCGACTG | CAGCTCGAGCTCGATGGATCCCTATCGGACGCCACCAAC |
| *Pto(MYB109)-LIKE* | GCCATGGAGGCCAGTGAATTCATGCAAAATCAAACTGAA | CAGCTCGAGCTCGATGGATCCTCAAGCAGGCATCTGGGT |
| *Pro-PtoSUS2-1* | GAAAAGCTTGAATTCGAGCTCGCAACGTACTTTTCAAGTGA | AGCACATGCCTCGAGGTCGACCTGGTGTTGATAATCAACCA |
| *Pro-PtoSUS2-2* | GAAAAGCTTGAATTCGAGCTCCATGGTTACTGAAGTGTTTGG | AGCACATGCCTCGAGGTCGACCGGTCTTAACCGTTTTATGC |
| *Pro-PtoVIN3-1* | GAAAAGCTTGAATTCGAGCTCCAACCAAATATTCAACTCACA | AGCACATGCCTCGAGGTCGACCAGCTTTGATGGGCCACG |
| *Pro-PtoVIN3-2* | GAAAAGCTTGAATTCGAGCTCGGTATGAACCTAAGATCACAC | AGCACATGCCTCGAGGTCGACGAGACCCATTTGCTAGGTAG |
